# EGFR/TBK1-dependent mitochondrial quality control contributes to acquired resistance to temozolomide

**DOI:** 10.64898/2025.12.19.695123

**Authors:** Lisenn Lalier, François Guillonneau, Abed-El-Razzak Saleh, Thomas Cabioch, Alice Boissard, Cécile Henry, Sophie Barillé-Nion, Judith Raimbourg, Philippe P. Juin

## Abstract

Acquisition of secondary resistance to chemotherapy in cancer cells is an ill-characterized multi-step process. We conducted a non-supervised, global proteomic analysis of U251 glioma cells treated by temozolomide as they transit from drug-sensitive to tolerant and then resistant states. This unprecedented analysis revealed key mitochondrial evolutions throughout the process. Following an early stress substantiated by a transient peak of reactive oxygen species, mitochondrial quality control mechanisms enable emergence of the resistant population. We combined genome editing, targeted protein degradation via PROTAC, and pharmacologic inhibitors to investigate the mechanisms underlying stress resolution both at population and single cell scales (including microscopic evaluation and *in-situ* proximity ligation assays). These studies unravel a Src-activated EGFR and TBK1 axis that initiates autophagic recycling of mitochondria as a necessary step to the emergence of the resistant population.

## Introduction

The adaptation capabilities of cancer cells significantly contribute to treatment-mediated acquisition of resistance. When submitted to cytotoxic chemotherapy, sensitive cancer cells undergo an initial stage of cell death, corresponding to a clinical improvement. However, even if disease then reaches a clinically undetectable level, some cancer cells often survive. The emergence of drug-resistant tumor populations is now conceived as resulting from a two-step transition, first described in lung cancer (1) and then identified in several types of cancers submitted to various therapies (2). In glioblastoma (GBM), the survival of a small fraction of the tumor cells population to the standard chemotherapy temozolomide (TMZ) was attributed to a stem-like phenotype (3), and the transition between the sensible to resistant populations was assigned to an epigenetic plasticity (2).

The transition from sensitive to resistant cells (2,4) first requires cells to evolve from a sensitive to a drug-tolerant stage, where cells survive despite slow cycling or the absence thereof. This stage clinically corresponds to a stable disease which corresponds to the minimal residual disease (MRD) and is under the control of both endogenous characteristics and microenvironmental factors. The duration of this stationary stage widely depends on tumor type, therapy and patients. Drug-tolerant cells (DTC) often exhibit increased disseminating abilities and are susceptible of migrating to distant niches, where they eventually participate to build a tumor-permissive microenvironment. The second major transition is from drug-tolerant to drug-resistant cells (DRC) which by definition differ from the DTC stage by being non reversible (that is, cells remain resistant even if drug is withdrawn). Moreover, DRC actively proliferate even in the presence of the drug and give rise to the disease recurrence, either in situ or in distant metastatic sites.

Mitochondria are complex integrators of cell death and survival signals. For instance, mitochondria are a major site of ROS production, through the mitochondrial respiratory chain which is required for energy production. These enzymatic reactions nevertheless produce ROS, which are detoxified by cellular Redox systems, preventing detrimental damages. In GBM, ROS play a bivalent role in the acquisition of resistance (5). Mitochondria are also central in the course of apoptosis via the control of the mitochondrial outer membrane permeability. Through communications with other intracellular membrane organelles, mitochondria also exchange various signals, regulating cell fate (6). These contacts also enable a dynamic shuttling of proteins from a compartment to another (7). Along with stress underwent by cells, especially cancer cells, the maintenance of healthy mitochondria integrity brings into play a process named mitochondria quality control (MQC) which involves dynamic mitochondria-associated membranes contacts, mitochondria fusion and fission processes and mitochondria recycling. Whether MQC is involved in the adaptation and resistance acquisition to TMZ, which players and pathways come into play, are still pending issues.

In this study, we took advantage of an in vitro model of resistance acquisition in GBM to explore the role played by mitochondrial adaptation, along with neighboring organelles interactions, in the emergence of the TMZ-resistant population. This model, first described in Rabé *et al.* (8), is based on the continuous treatment of U251 cells by a clinically relevant dose of temozolomide (TMZ). This model is highly reproducible with a well-defined chronology and enables the emergence of the resistant population with a proliferation rate similar to the naive population within about three weeks. We performed a non-supervised proteomic analysis of kinetic changes in the proteome of U251 cells acquiring resistance to TMZ. This analysis hinted on a major role played by mitochondrial adaptation during this process. Further analysis shed the light on a new role played by the couple EGFR-TBK1 in the MQC and the resolution of stress required for the transition of cells from the DTC population to the DRC population.

## Materials and methods

### Antibodies and drugs

#### Drugs

TMZ (MedChemExpress #HY-17364); NAC (Sigma Aldrich #7250); PP1 (Sigma Aldrich #P0040); saracatinib (Interchim #11497); erlotinib (Interchim #10483); 3MA (Euromedex #S2767).

#### Antibodies

TOM20 #612278, BD (Le Pont de Claix, France); TFAM #8076, Cell Signaling Technology (Danvers, MA, USA); EGFR #4267, Cell Signaling Technology; pEGFR Y845 #6963, Cell Signaling Technology; pEGFR Y1068 #3777, Cell Signaling Technology; TBK1 #3504, Cell Signaling Technology; pTBK1 #5483, Cell Signaling Technology; pSrc #sc-81521, Santa Cruz Biotechnology (Dallas, TX, France); VDAC #ab14734 Abcam, (Cambridge, UK); IP3R #sc-28614, Santa Cruz Biotechnology; LC3 #83506, Cell Signaling Technology or #L7543, Sigma Aldrich.

#### Other reagents

MitoSOX Red #M36008, Invitrogen (Thermo Fisher Scientific, Waltham, MA, USA); Rhod2-AM #R1244, Invitrogen; Prolong Gold with DAPI #836941, Invitrogen; PROTAC TBK1 and negative control TBK1 PROTAC 3i (#7259) and TBK1 control PROTAC 4 (#7260), Tocris Bioscience (BioTechne SAS, Noyal Chatillon sur Seiche, France).

### Statistics

Unless otherwise stated, experiments were performed in triplicates and data were analyzed by GraphPad Prism by unpaired Student t-test.

### Cells culture and modifications

U251 cells were purchased from the ATCC. They were grown in DMEM medium (4.5g/l glucose) supplemented with 10% FBS, glutamine 2mM and penicillin/streptomycin. Cells were treated by the indicated drugs as follows: TMZ 50µM, NAC 5mM, PP1 10µM, 3-MA 5mM, TBK1 or control PROTAC 100nM.

U251 Rho0 (ρ^0^) cells were obtained by the constant culture in defined medium containing 50 ng/mL EtBr (ethidium bromide), 50 µg/mL uridine and 110 ng/mL pyruvate (ρ^0^ medium) for at least 4 weeks before testing to determine whether the cells exhibited ρ^0^ -like mitochondrial damage (altered mitochondrial morphology, inability to grow without uridine/pyruvate supplementation).

U251 EGFR-cells were obtained by the transfection of a set of two plasmids purchased from LabOmics (Belgium), one expressing Cas9 and sgRNA targeting EGFR gene (#HCP204605-CG01-3-B) and one genome-editing donor conferring resistance to puromycin (#DC-HTN204605-D07). Cells were transfected by the use of JetPrime reagent (Polyplus) according to the manufacturer instructions and were further selected by puromycin 1µg/ml.

mEmerald-Mito-7 was a gift from Michael Davidson (Addgene plasmid # 54160). U251 cells were transfected by the use of JetPrime reagent (Polyplus) according to the manufacturer instructions and were further selected by G418 (1mg/ml). To ensure optimal expression, cells were sorted with a BD FACSAria III cytometer by the Cytocell core facility.

### Cell count for kinetics experiments

U251 cells were seeded in 24-well plates. TMZ 50µM was added twice a week. At the indicated time points, cells were trypsinized, pelleted and resuspended in PBS. Cell count was determined by flow cytometry (Accuri C6 or Attune Nxt cytometer) and normalized by the number of cells counted at D0. Data were obtained from four replicates.

### Proteomics experiments (performed at the Prot’ICO proteomics facility)

Sample preparation: The pelleted cells (300 thousands cells, sorted from 5 independent biological replicates per conditions) from each time point were quickly thawed and proteins were concomitantly extracted and denatured using 200µL of 0.1% Rapigest-SF® (Waters), 5mM DTT and 50mM TRIS pH 8.5, at 95°C for 30min. Sonication was performed 30s at 20% power using a probe sonicator. Thiol residues were thus chemically reduced (by DTT) and subsequently protected by alkylation for 10min at 37°C in 10mM MMTS (Sigma). Samples were cooled to room temperature before adding 2µg trypsin (Porcine, TPCK treated, from ABSciex), incubated at 37°C overnight with 750rpm mixing on a thermomix. Resulting peptides were then acidified using 23µL of Formic Acid (FA) for one hour at 37°C under 750rpm thermomixing to break down Rapigest-SF® and insoluble material was cleared by centrifugation. The supernatant-containing peptides was desalted using in-house pipette tips-packed with C18 described as “Stage-tips” procedure (9) and eluted. Eluates were dried in a vacuum centrifuge concentrator (Thermo), resuspended in 0.1% FA. The equivalent of 200ng of peptides were injected in LC-MS/MS after a microBCA™ peptide assay (Thermo Fisher Scientific) using the Hystar hardware management software suite v. 6.3.1.8 (Bruker).

Ultra-Performance Liquid Chromatography (LC): Each sample was injected and separated on a C18 reverse phase column (Aurora ultimate series 1.7µm particles size, 75µm inner diameter and 25cm length from IonOptics) using a NanoElute LC system (Bruker). Eluate flow was electrosprayed into a timsTOF Pro 2 mass spectrometer (Bruker) for the 60min duration of the hydrophobicity gradient ranging from 99% of solvent A containing 0.1% FA in milliQ-grade H2O to 40% of solvent B containing 80% ACN plus 0.1% FA in mQ-H2O.

Mass Spectrometry (MS) data acquisition: The mass spectrometer operated throughout the elution process in Data-Independent Analysis (DIA) mode, with PASEF-enabled method using the TIMS-Control software 5.1.8 (Bruker). The m/z acquisition range was 300–1201 with an ion mobility range of 0.75–1.25 1/K0 [V s/cm2], which corresponded to an estimated cycle time of 1.06 seconds at 100% duty cycle. DIA-PASEF windows, and collision energy were also left to default with a base of 0.85 1/K0 [V s/cm2] set at 20eV and 1.3 1/K0 [V s/cm2] set at 59eV. TOF mass calibration and TIMS ion mobility 1/K0 were performed linearly using three reference ions at 622, 922, and 1222 m/z (Agilent Technology, Santa Clara, CA, USA)

LC-MS raw data analysis, evaluation, protein identification and quantification: The raw data was generated by DIA-PASEF. Identification and quantification data were extracted using Spectronaut® v.19.0.240606.62635. In short: DirectDIA+ (library-free) mode allowed generation of modelized peptides out of the human UniProt Knowledge Base of model organism_9606 (version 2024, april 3^rd^ with 42 514 entries) in FASTA format. The enzyme’s cleavage specificity was that of trypsin’s. The precursors and fragments’ mass tolerances were set to a maximum of +/-15ppm variations. Modification of methionines by oxidation and acetylation of protein N-termini were set as possible events while all thiol groups from cysteines were considered completely alkylated by methylation. The software’s deep learning algorithm permitted an overall range of identification of over 8000 protein groups. Data were normalized between samples, match between runs was allowed to interpolate missing identifications. No data imputation was performed for ratio calculations. A threshold of adjusted p-value <0.01 was applied so the output (precursors) was filtered to maintain false discovery rate below 1%. Retention time alignment and correction for mass accuracy were performed automatically. Spectronaut® statistical tools were used to visualize data, assess performances and quality before quantitatively compare groups of 5 replicates’ datasets. Unsupervised principal component analysis of the whole dataset is displayed in suppl fig1b. The resulting proteins LFQ values were log2(x) transformed to express the relative abundance of proteins. We defined thresholds for up- and down-regulated proteins as absolute (log2Ratio) >= 0.58 and adjusted p-value (or q-value) < 0.05. These ratios were defined to determine the level of modification of the protein expression.

The lists of candidates significantly different between two time points identified as described above, either up- or down-regulated, were submitted to GSEA analysis via Enrichr. GSEA lists used are Gene Ontology Biological Process 2023 and Cellular Component 2023. The lists of proteins upregulated at D6 vs D0 and downregulated at later time points were superimposed and represented by Venn diagrams. The list of common proteins was then submitted to GSEA analysis as described above. Prism software was used to draw graphs from the data obtained by Enrichr and for statistical analyses.

### Immunocytofluorescence

Cells were seeded at D-1 on glass coverslips and eventually treated with the indicated drugs from D0 for the indicated time. Cells were rinsed by PBS, fixed by PBS-4% paraformaldehyde for 15 min at room temperature and kept in PBS at 4°C. Cells were permeabilized by Triton X100 0.5% and saturated by PBS-5% BSA. Primary and AF647-coupled secondary antibodies were incubated in PBS-1% BSA1%-0.1% saponin. Coverslips were mounted in Prolong Gold medium with DAPI for nucleus staining. Image acquisition was performed with a Nikon A1 Rsi confocal microscope. Data were analyzed by the ImageJ 1.54g software. The analysis of mitochondrial morphology was performed by Mitochondria Analyzer (Fiji plugin) according to the manual after image 3D deconvolution performed by NIS software (block size 1.35, Cvalue 11). Each image was analyzed as a whole and the number of mitochondria was divided by the number of DAPI-stained nuclei to obtain the mean number of mitochondria per cell.

### Flow cytometry

Flow cytometry was performed with BD Accuri C6 or ThermoFisher Attune Nxt cytometer. For the detection of intracellular proteins, cells were fixed by 4% paraformaldehyde, permeabilized by 0.5% saponin and saturated by 5% BSA. Antibodies were diluted in PBS-0.1% saponin-1% BSA and coupled to fluorescent (AF647) secondary antibodies. MFI were corrected by the fluorescence measured by cells incubated with the secondary antibody only.

### Proximity Ligation Assay (PLA)

Cells were seeded in 24-well plates onto glass sterile coverslips and allowed to attach overnight. The indicated treatment was applied, then cells were fixed by 4% paraformaldehyde. The labeling with antibodies was realized according to the manufacturer instructions (Duolink, Sigma-Aldrich). Coverslips from each kinetic experiment were treated simultaneously as paired samples and the mean number of spots per cell at each timepoint was normalized to the related number of spots at D0. Negative controls consist of coverslips treated as the others except from primary antibodies omission. Images acquisition was performed with a Nikon A1 Rsi confocal microscope. Data were analyzed by the ImageJ 1.54g software and spots were quantified by the Object 3D Counter (Fiji plugin).

### MitoSOX and Rhod2AM staining

Cells were seeded in 6-well plates and treated as indicated. Cells were stained by MitoSOX 1µM for 20min or Rhod2AM 1µM for 15min. Cells were rinsed and harvested in PBS before fluorescence measurement by flow cytometry.

### IC50 measurements

IC50 measurements were measured as described earlier (10) by the MTT method. Briefly, cells were plated in 96-well plates (3000 cells/well) and treated by the indicated drugs. Cells were incubated with MTT 3h at 37°C, formazan cristae were solubilized in DMSO and OD was read at 570nm. IC50 was calculated by the AAT Bioquest “IC50 Calculator” tool (AAT Bioquest, Inc. (2025, February 17). *Quest Graph™ IC50 Calculator*. AAT Bioquest. https://www.aatbio.com/tools/ic50-calculator)

## Results

### TMZ induces mitochondrial modifications during the process of resistance acquisition in U251 cells

When submitted to a constant treatment by TMZ at a clinically relevant dose, U251 cells undergo successive stages as described by Rabé *et al.* (8) (figure 1A). Consistent with previously published results, maximal cell death rates occur after three to six days TMZ-treatment (D3-D6) and later regress (suppl fig 1a). The cell population size is minimal around D6 to D9, then cells progressively begin to proliferate and give rise to the so-called resistant population around D21, which exhibits a rate of proliferation similar to parental cells (figure 1A).

**Figure 1:**
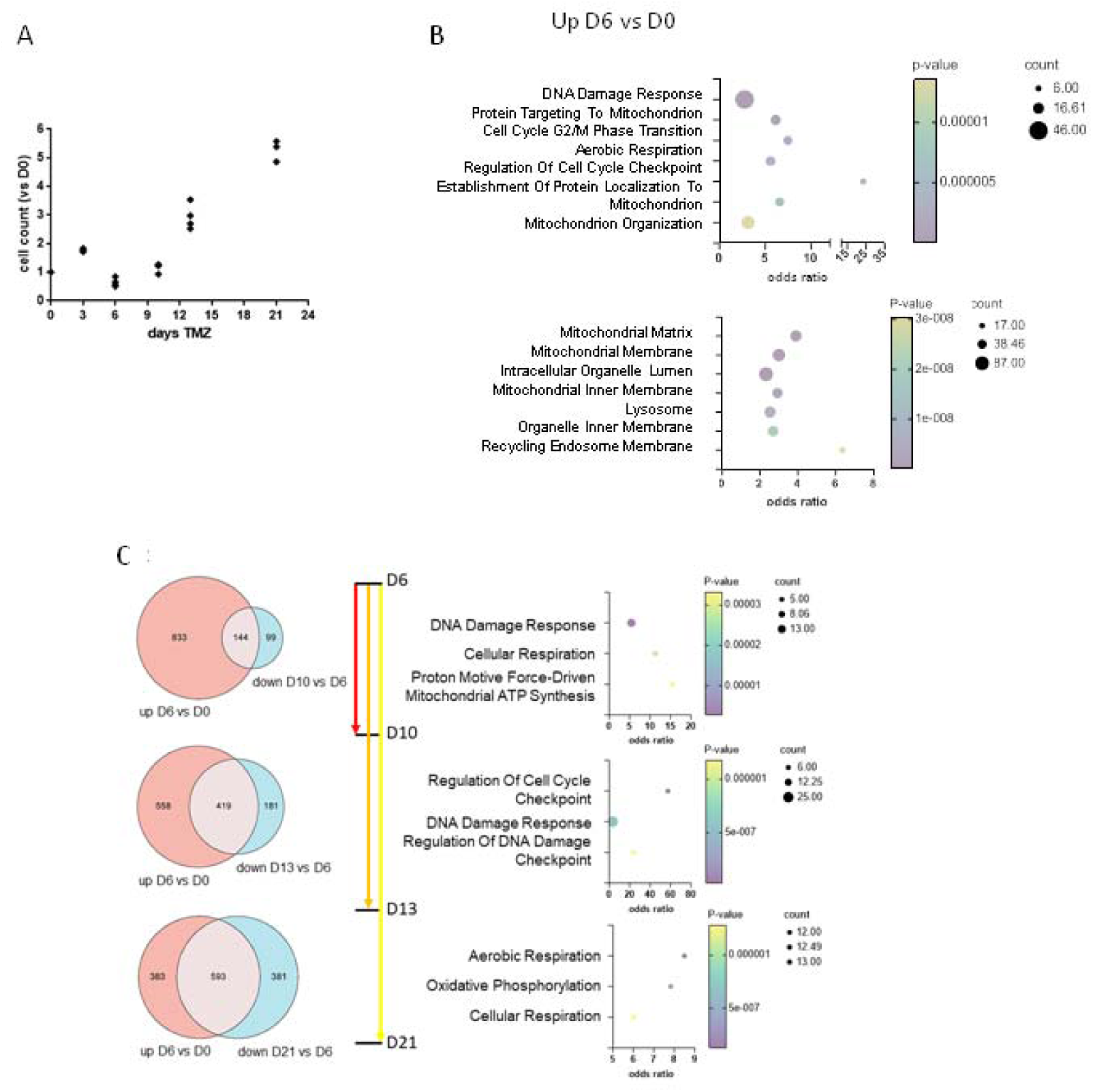
A. Evolution of the U251 cell population during TMZ treatment. U251 cells were treated by TMZ 50µM every 3-4 other day and cells were counted by flow cytometry at each time point. Cell number was normalized by the cell count at D0. B. Proteins significantly overexpressed at D6 vs D0 in the 5 mass spectrometry experiments were listed and submitted to GSEA analysis (Enrichr). The 7 top pathways identified by Gene Ontology Biological Process (up) and Gene Ontology Cell Component (bottom) are represented on the bubble graphs. C. The overlap between proteins overexpressed at D6 vs D0 and down-regulated at later time points (D10, D13 and D21 respectively) vs D6 is represented on the Venn diagrams (left). The list of proteins present in the intersection was submitted to GSEA (Enrichr) and the 3 top pathways identified by Gene Ontology Biological Process are represented on the bubble graphs (right).

We explored the global proteome of U251 cells throughout the process of TMZ-resistance acquisition by mass spectrometric analysis of TMZ-treated cells at six time points: D0 before the treatment initiation, D3 at the beginning of detected cell death, D6 when cell death declines, D10 when the first proliferation foci are seen, D13 when cells form an intermediate population with an active proliferation and D21 when cells regain a similar proliferation rate to parental cells.

The analysis was conducted on five sample replicates, obtained from independent resistance acquisition experiments. Proteomic analyses identified more than 8000 protein groups (more than 12000 proteins) in every sample. As revealed by a non-supervised principal component analysis (PCA) (suppl fig 1b) the different replicates of the same time points tended to segregate together, although not for all of them. This illustrates the increase of the relative phenotypic heterogeneity between replicates in the intermediate time points (D3, D10 and D13) that partly overlap each other, whereas D0, D6 and D21 constitute much better segregated clusters. Of note, the second dimension of PCA segregates D0 from D21 cells whereas they are close on the first dimension. This PCA illustrates that D6 cells are extremely different from D0 and D21 cells, which is consistent with the notion that their phenotype represents a peak of cell stress. A gene set enrichment analysis (GSEA) performed by Enrichr (11) highlighted the biological pathways either up- or down-regulated between two time points. The top biological processes up-regulated at D6 vs D0 (figure 1B and suppl fig 1c) include DNA damage response and regulation of cell cycle, which most likely relate to the initial toxicity to DNA induced by TMZ. The second top cluster of biological processes up-regulated at D6 reveals the mitochondrial adaptation of cells, including aerobic respiration, emphasized by the predominance of the mitochondrial compartments as the top cellular components upregulated at D6. Meanwhile, biogenesis is globally inhibited, probably linked to the reduced cell proliferation. We then questioned the process of stress resolution form D6 to D21 at the gene set enrichment level (figure 1C). The direct comparison of D0 vs D21 cells shows a persisting metabolic modification in D21 cells, which are predicted to rely less on mitochondrial respiration than parental cells, and express more proteins related to cytoskeleton modifications linked to mitosis (suppl fig 1d). Of note, the upregulation of pathways associated to cell division at D21 (suppl fig 1d) may be associated to the passage of cells we had to perform the day before collection for mass spectrometric analyses. From the peak of stress at D6 and then on, proteomic analysis suggests that the population tends to evolve toward a population more and more similar to the initial one, even if not identical (figure 1C, circles). The overlap of proteins overexpressed at D6 vs D0 and proteins down-regulated at later time points compared to D6 highlights the progressive resolution of the modifications induced at D6. This resolution towards the resistant population at D21 mostly concerns the mitochondrial oxidative phosphorylation process, upregulated at D6, and the protein synthesis, repressed at D6 (suppl fig 1e). From D6 to D10, during the transitory stage, cells progressively overcome DNA damage, which appears as the highest-scored pathway (figure 1C, up). This resolution is amplified at D13, as illustrated by the Venn diagram (figure 1C, middle), while pathways associated to DNA repair, mitosis and transcription are highlighted. Strikingly, pathways associated to the mitochondrial metabolic adaptation are the second cluster of top pathways identified by the gene set enrichment analysis and the top clusters of the overlap between proteins overexpressed at D6 vs D0 and downregulated at D21 vs D6 (figure 1C, down). These data prompted us to further investigate the mitochondrial evolution along the process of resistance acquisition.

### Mitochondrial morphology transiently evolves along with mitochondrial transcription

First of all, we sought for a possible correlation between the evolution of mitochondria during TMZ treatment and the evolution of the cells proteome we have identified. At the morphological level, as evaluated by fluorescent microscopy, the mitochondrial network acquired a fused, tubular phenotype during the drug-tolerant stage, obviously observed at D9 (figure 2A and suppl fig 2a), with a progressive increase in the mitochondrial mean volume (figure 2B). At day 12, when cells progressively regain their proliferation capacity, mitochondria divided and became smaller, resulting in a transient significant increase in the mean number of mitochondria per cell (figure 2B). At the proteomic level, grouped fold change analysis highlighted a few proteins implicated in the mitochondrial dynamics differentially expressed during the acquisition of resistance. We calculated the expression fold change (log2FC) of each protein vs its paired expression at D0 as represented on suppl fig 2b. Surprisingly, proteins involved in the mitochondrial fusion process were not significantly upregulated in the intermediate time points. We could only notice a relative increase after D6 in OMA1 (D13 vs D6 p=0.0095) and in MFN1 (D10 vs D6 p=0.0074 and D13 vs D6 p=0.0051). The morphological change of the mitochondrial network might thus be rather attributed to a decrease in fission proteins before D6 (MTFRL1 D3 vs D0 p=0.0472 and D6 vs D0 p=0.0122, MIEF1 D6 vs D0 p=0.015, FIS1 D3 vs D0 p=0.0378) before they eventually raise (MTFRL1 D13 vs D6 p=0.017). SLC25A46 is significantly overexpressed at the time points when mitochondrial fragmentation is observed (D10 vs D0 p= 0.0311 and D13 vs D0 p=0.038) whereas MTFR2 is overexpressed at all intermediate time points (p=0.159 at D3, p=0.004 at D6, p=0.025 at D10, p=0.0196 at D13 vs D0 respectively). However, it should be noted that MTFR2 was detected at a modest level throughout the kinetics (max quantity = 61.27). Finally, MFF isoform 2 was detected at the maximal level at D6 and the expression progressively decreased relative to D6 (D10 vs D6 p=0.0141, D13 vs D6 p=0.0012).

**Figure 2:**
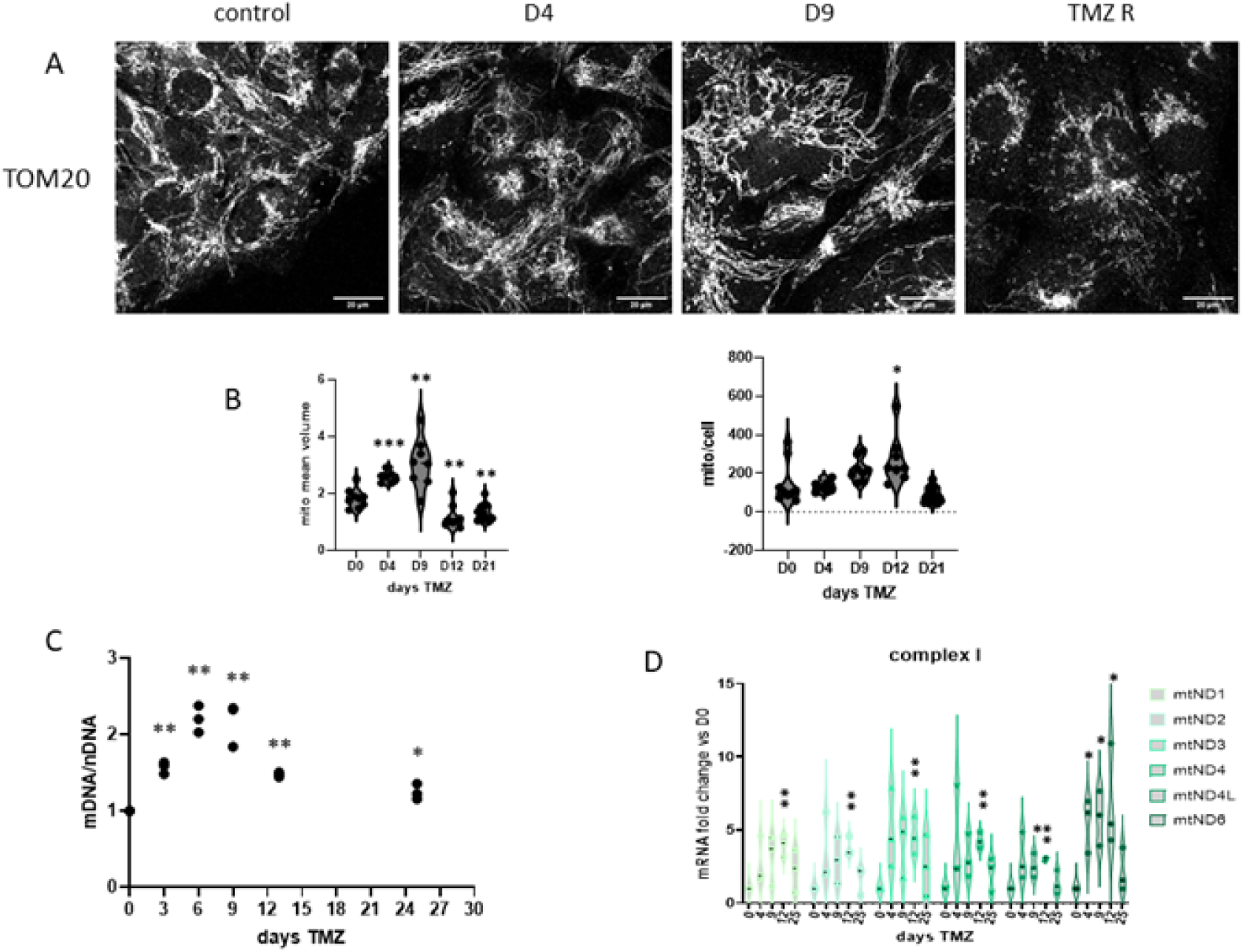
A. Mitochondria from U251 cells treated by TMZ during the indicated time were stained by anti-TOM20 antibody and observed by confocal microscopy. The images shown are representative from three independent experiments (scale bar: 20µm). B. The mean mitochondria volume and number of mitochondria was estimated by the Fiji software (Mitochondria Analyzer plugin) and divided by the number of cells. The analysis was performed from at least six images per time point. (*: p<0.05, **: p<0.01, ***: p<0.001 vs D0). C. Total DNA was extracted from TMZ-treated U251 cells and the ratio mtDNA/nDNA was calculated as described. The graph illustrates three independent experiments (**: p<0.01, ***: p<0.001 vs D0). D. RT-QPCR was performed on total mRNA extracted from TMZ-treated U251 cells. RPLPO was used as housekeeping gene and non-treated U251 cells were used as reference. The graph illustrates data from three independent experiments (*: p<0.05, **: p<0.01 vs D0).

Since mitochondrial morphology parallels mtDNA replication, we measured the ratio of mtDNA/nDNA during the process of resistance acquisition as described in Moretton *et al* (12). As indicated by figure 2C, the mtDNA/nDNA ratio rapidly increased when TMZ was added to reach a maximum in TMZ-tolerant cells (D6 to D9) and then progressively returned to the initial level in TMZ-resistant cells. We then measured the transcription of the mRNA of 12 out of 13 mitochondria-encoded proteins. As shown by figure 2D and suppl fig 2c-d, this increase in mtDNA replication was followed by an increase in the translation of mitochondria-encoded mRNAs, mostly significant at D12, with some variability between mRNAs. In the proteomic analysis, among the candidates differentially expressed associated to mtDNA replication according to Human MitoCarta3.0 (suppl fig 2e), several proteins were significantly overexpressed at the intermediate time points vs D0 (POLG2, SSBP1, TFAM, TOP1MT, MGME1, paired Student t-test). ENDOG was the most downregulated protein, the only significant difference being D6 vs D0 (mean fold change 0.24 p=0.0366). DNA2 and LIG3 were very significantly downregulated at D6 vs D0 (p<0.0001) and the decrease remained significant respectively at D10 and D10 and D13. Of note, TFAM, TFB2M and TOP1MT were associated to both mtDNA replication and transcription according to Human MitoCarta3.0. The proteins TEFM and MRPL12 were also downregulated at D6 vs D0 (p=0.0356 and 0.0063 respectively), while MRPL12 was significantly downregulated from D3 to D10. Of note, despite an overt global increase in mRNA levels of mitochondria-encoded proteins, overexpression of the corresponding proteins was not detectable by proteomic analysis.

### Oxidative metabolism is increased during the intermediate stages of resistance acquisition

GSEA analysis highlighted aerobic respiration increase during the intermediate time points of the resistance acquisition (figure 1). We therefore questioned the expression of proteins from the respiratory complexes (complex I to IV, thereafter designed CI to CIV) in the proteomic analysis. Although proteins from the CII were globally overexpressed during the transitory stage, the expression of proteins from the other complexes was less consistent, some proteins being overexpressed and other being downregulated (suppl fig3a). Nevertheless, mtROS (evaluated by MitoSOX staining) actually peaked at D6 and then rapidly decreased (figure 3A). This peak in mtROS production happened together with an increase in the mitochondrial Ca2+ concentration, as measured by the fluorescent dye Rhod2AM (figure 3B). The mitochondrial localization of the Rhod2AM staining was confirmed by confocal microscopy (suppl fig 3b) performed on U251 mEmMito7 cells, expressing a fluorescent mitochondrial protein (mEmerald-Mito7, see material and methods section). ER is the major regulator of the cytoplasmic regulation of Ca2+ concentration, confining Ca2+ in ER lumen. Ca2+ transfer from the ER to the mitochondria happens through the close apposition between IP3R at the ER membrane, VDAC1 at the mitochondrial outer membrane (MOM), stabilized by Grp75, and MCU at the mitochondrial inner membrane (MIM), thereby enabling the entry of ER Ca2+ into the mitochondrial matrix (13). As generally observed during stress (14,15,16), an increase in ER-mitochondria contacts was detected as an early reaction to TMZ with a peak at D6, objectified by the amount of VDAC-IP3R proximity ligation assay (PLA) dots (suppl fig 3b), with a progressive return to the initial level (figure 3C and suppl fig 3c). MCU was overexpressed from D3 to D10 vs D0 (p=0.017 at D3 and p<0.001 at D6 and D10), whereas MICU1 was overexpressed at D13 vs D0 (p=0.048). Simultaneously, EFHD1, negative regulator of Ca2+ mitochondrial uptake, was downregulated from D6 to D13 vs D0 (p<0.01). By contrast, RHOT1 was significantly downregulated at D6 vs D0 (p<0.01). Interestingly, the Ca2+ transporter LETM1, which also plays a role in the mitochondria and cristae structure and in the assembly of respiratory complexes, was significantly overexpressed at the intermediate time points (p<0.01 at D3, D6 and D13 vs D0, p<0.0001 at D10 vs D0) (figure 3D).

**Figure 3:**
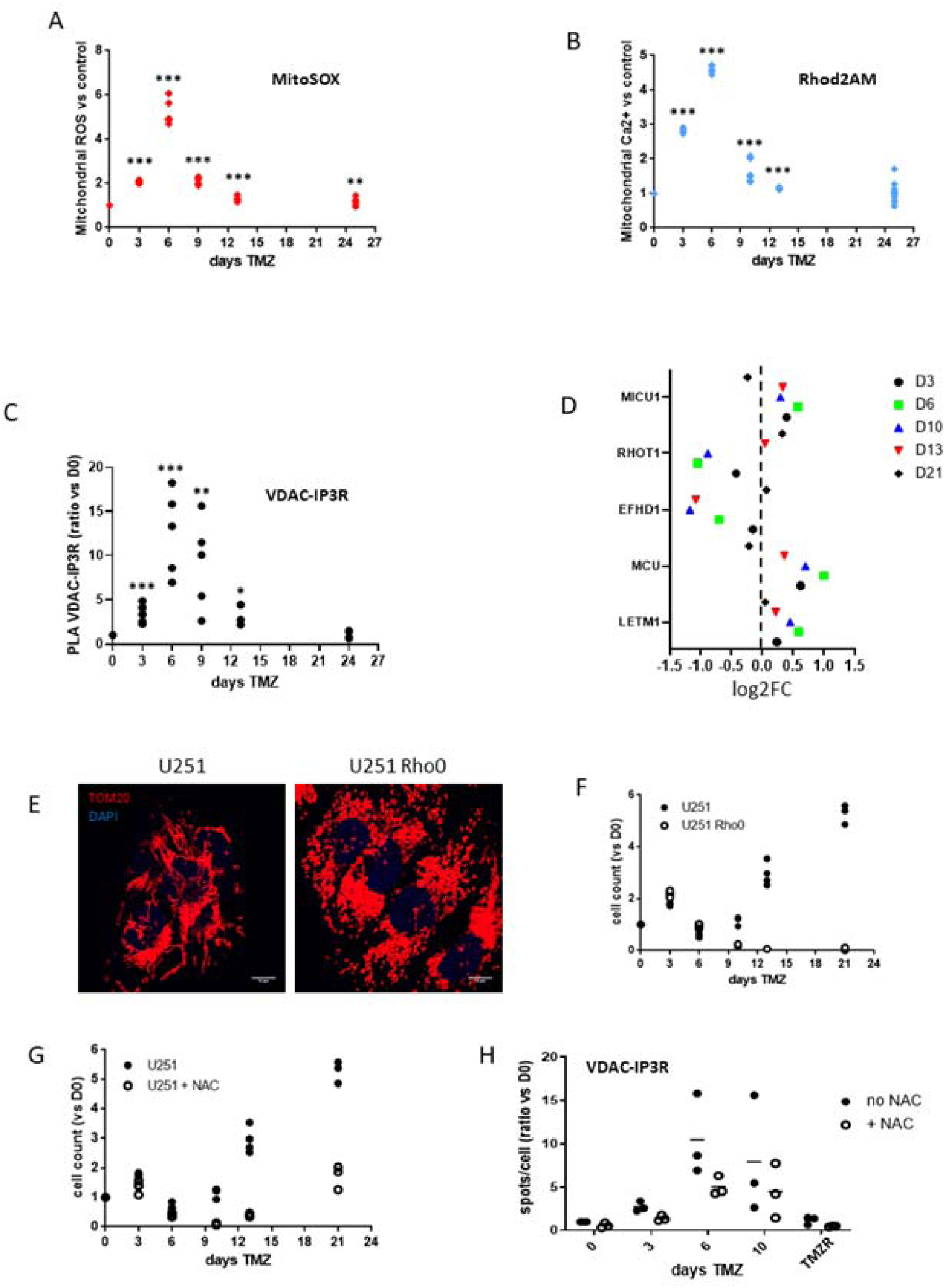
U251 cells were treated by TMZ for the indicated time and stained by MitoSOX (A) or Rhod2AM (B). The mean fluorescence was normalized by the mean fluorescence of non-treated U251 cells. Data shown are replicates of at least six measurements (**: p<0.01, ***: p<0.001 vs D0). C. VDAC-IP3R PLA was performed on TMZ-treated U251 cells. The number of spots was divided by the number of cells on each image and normalized by the number of spots per cell at D0. (*: p<0.05, **: p<0.01, ***: p<0.001 vs D0). D. The log2FC values of candidates differentially expressed and involved in Ca2+ signaling (MitoCarta) were calculated for each of the five replicates and the mean of the five values is represented on the graph. The results of the paired Student t-tests are described in the main text. E. U251 and U251 Rho0 cells were fixed and stained with TOM20 antibody and DAPI. The images shown are representative of three independent experiments (scale bar: 10µm). F. U251 and U251 Rho0 cells were treated by TMZ 50µM every 3-4 other day and cells were counted by flow cytometry at each time point. Cell number was normalized by the cell count at D0. G. U251 cells were treated and counted as in F with or without NAC 5mM added to each TMZ treatment. H. VDAC-IP3R PLA was performed as in C. with or without NAC 5mM added to each TMZ treatment.

To further explore the importance of mitochondrial activity in the acquisition of TMZ resistance, we investigated the long-term behavior of respiratory deficient U251 Rho0 cells. To this purpose, we generated mtDNA deficient U251 Rho0 cells by the long-term culture of cells in the presence of EthBr (50ng/ml). These cells require a supplementation in uridine and pyruvate to be able to grow and proliferate, although at a slower rate than U251 cells. The Rho0 cells exhibit non-functional, round-shaped mitochondria (figure 3E) and produce no mtROS, either at the basal level or when treated by TMZ for 72h (suppl fig 3d). When treated by TMZ, U251 Rho0 cells first behaved similarly to U251 cells (even if the Rho0 cells appeared initially less sensitive to TMZ-induced cell death), but they proved unable to escape from the TMZ tolerance and no viable cells were counted after 10 days of culture (figure 3F). As we previously observed an increase in mtROS production in TMZ-treated U251 cells, we added NAC 5mM to each TMZ treatment to explore the consequence of dampening their accumulation. Of note, probably owing to the short half-life time of NAC, this treatment did not totally preclude the fluctuation in mtROS level. However, as indicated in figure 3G, NAC significantly delayed the emergence of the TMZ-resistant U251 cells, revealing the hormetic property of the ROS early peak observed. Furthermore, NAC reduced the peak of VDAC-IP3R PLA spots by about 40% at every time point (figure 3H and suppl fig3e), suggesting that ROS regulate ER-mitochondria contacts during TMZ treatment, which in turn may feedback amplify ROS production. The afflux of Ca2+ via ER-mitochondria contacts is namely an activator of the mitochondrial oxidative activity, resulting in ROS production (17). These results argue for an orchestrated reciprocal regulation between mitochondrial-ER contacts and mitochondrial oxidative activity in the acquisition of TMZ resistance.

### TBK1 phosphorylation induced by TMZ requires Src-activated EGFR, which is necessary to the emergence of the TMZ-resistant population

The mtDNA deficiency of respiratory disabled Rho0 cells prevents them from signalling through mtDNA cytoplasmic release, which in particular activates the cytoplasmic DNA sensing cGAS-STING axis upstream of TBK1. We observed the activation of TBK1 at D6 and D9 during TMZ treatment (figure 4A). However, the knock-down of either cGAS or STING (18) did not modify the kinetics of resistance acquisition to TMZ or the phosphorylation of TBK1 (suppl fig 4a, b). This indicates that a cGAS/STING independent (and most likely mtDNA release independent) TBK1 activating mechanism occurs during resistance acquisition.

**Figure 4:**
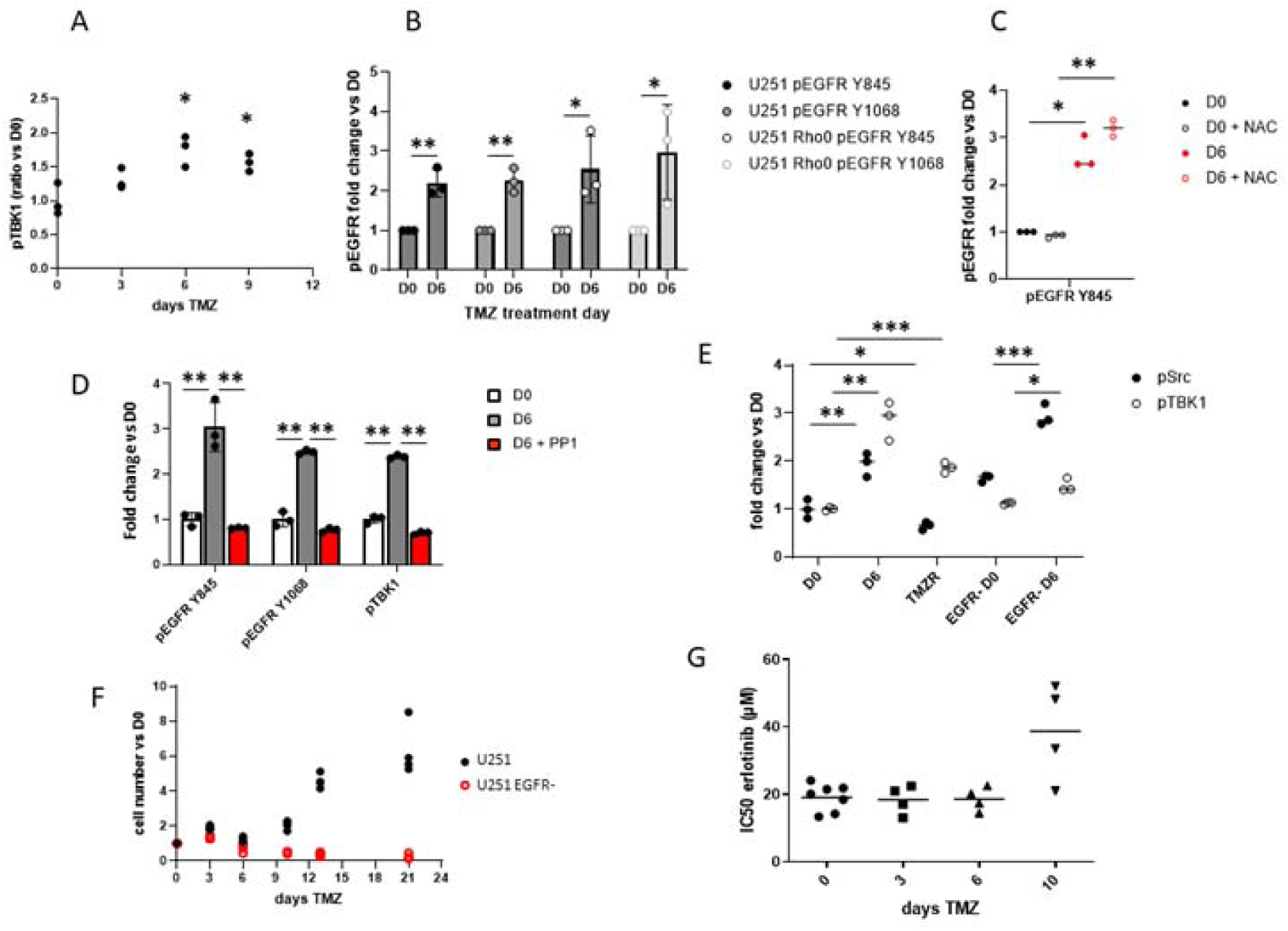
A. U251 cells were treated by TMZ. Cells were harvested, fixed and stained by pTBK1 antibody coupled to fluorescent secondary antibody. pTBK1 specific fluorescence was detected by flow cytometry and expressed as a ratio to D0 fluorescence (*: p<0.05 vs D0). B. EGFR phosphorylation was measured by flow cytometry by the use of phospho-specific antibodies (pEGFR Y845 or pEGFR Y1068) on U251 or U251 Rho0 cells. Fluorescence was normalized to the fluorescence measured at D0 (*: p<0.05, **: p<0.01). C. EGFR Y845 as measured as in B. in the presence or absence of NAC 5mM (*: p<0.05, **: p<0.01). D. Phosphorylation of EGFR (Y845 and Y1068) and TBK1 was measured as in B in the presence or absence of PP1 (Src inhibitor). E. Src and TBK1 phosphorylation was measured as in B in U251 and U251 EGFR-cells treated by TMZ for the time indicated on the graph (*: p<0.05, **: p<0.01, ***: p<0.001). F. U251 and U251 EGFR-cells were treated by TMZ 50µM twice a week and cells were counted by flow cytometry at each time point. Cell number was normalized by the cell count at D0. G. The sensitivity of U251 cells to erlotinib was evaluated by MTT during TMZ treatment and the IC50 was calculated by AAT Bioquest “IC50 Calculator” tool (https://www.aatbio.com/tools/ic50-calculator).

As an alternative pathway for TBK1 activation, Chakraborty *et al.* (19) demonstrated in U251 cells that the non-canonical, ligand-independent activation of EGFR induces the recruitment and consecutive activation of TBK1. We have previously demonstrated in lung cancer that chemotherapy induces EGFR non canonical activation in a ligand-independent, Src-dependent way (10). We therefore decided to further investigate the role of Src-activated EGFR in TMZ resistance. At the beginning of the TMZ-tolerant stage (D6), we measured a significant phosphorylation of EGFR Y845, the target of Src responsible for its mitochondrial targeting (20), and of EGFR Y1068, a marker of EGFR kinase activation (figure 4B). Of note, the increase in EGFR phosphorylation was similar in U251 Rho0 cells, suggesting that mtROS is not the initiator of EGFR phosphorylation. Similarly, NAC did not reduce the phosphorylation of EGFR Y845 (figure 4C). By contrast, the inhibition of Src kinase activity by PP1 or saracatinib (suppl fig 4c) abolished both EGFR Y845 and EGFR Y1068 phosphorylation (figure 4D) without reducing total EGFR expression (suppl fig 4d). We also noticed that TBK1 phosphorylation at D6 is inhibited by PP1 (figure 4D). To reveal the role played by EGFR, we knocked-out EGFR in U251 cells (without altering cell viability) and treated them by TMZ. The absence of EGFR did not prevent the activation of Src (figure 4E), whereas it drastically reduced the activation of TBK1 (2.9-fold increase in U251 cells vs 1.3 in U251 EGFR-). When treated by TMZ for a longer period, U251 EGFR-cells did not recover their proliferative activity and the population progressively died (figure 4F). We have previously observed in NSCLC that EGFR non canonical activation by cisplatin induced a sensitization to EGFR tyrosine kinase activity inhibition by erlotinib (10). We thus questioned the role played by EGFR tyrosine kinase activity in the present resistance process by treating U251 cells with erlotinib during the acquisition of TMZ resistance. We measured no alteration of the IC50 to erlotinib (72 hour-treatment) from D0 to D6 of TMZ treatment, rather a tendency to increase the IC50 at D10 (figure 4G), arguing that EGFR mitochondrial localization rather than its kinase activity is required for U251 cells to acquire the resistance to TMZ.

### Src-activated EGFR is involved in the early modifications of the mitochondrial network but also in the recovery phase

We investigated the effect of Src inhibition via PP1 before (D0-D6) or after (D6-D13) the ROS peak. Surprisingly, PP1 prevented the recovery of U251 cells proliferation both when cells were treated before or after the peak of ROS (figure 5A). Although more toxic than PP1 on U251 cells, saracatinib (a dual inhibitor of Src and Bcr-Abl kinases) efficiently inhibited Src activation induced by TMZ (suppl fig 4a) and the number of viable cells when added before or after D6 (suppl fig 5a), even if the effect was less drastic than PP1 in the conditions we used. At the morphological level, U251 EGFR-cells did not exhibit the same evolution as U251 cells when treated by TMZ (Figure 5B). We detected no shift of mitochondria to a tubular, fused network, as mitochondria instead tended to be more fragmented (figure 5B, C). We neither observed any increase in ER-mitochondria contacts in U251 EGFR-cells (suppl fig 5b). The number of mitochondria per cell tended to increase slightly even if the difference between D0 and D9 was not significant (figure 5D). Meanwhile, the amount of mtDNA did not significantly increase (figure 5E) and the translation of mitochondria-encoded mRNA did not significantly increase (suppl fig 5c). Of note, the expression (suppl fig 2e) and distribution pattern of TFAM (suppl fig 5d), the major factor governing mtDNA compaction and translation, seemed to evolve during TMZ treatment in U251 cells, possibly contributing to the modulation of mtDNA expression observed above (figure 2c). By contrast, in EGFR-deficient cells, TFAM exhibited the same expression pattern from D0 to D9 (suppl fig 5d). This phenomenon probably ensues from an indirect relationship however since the mitochondrial membrane in resting U251 cells is not the mainstream localization of EGFR. Altogether these results suggest a role played by Src-activated EGFR both before the ROS peak and during the recovery phase.

**Figure 5:**
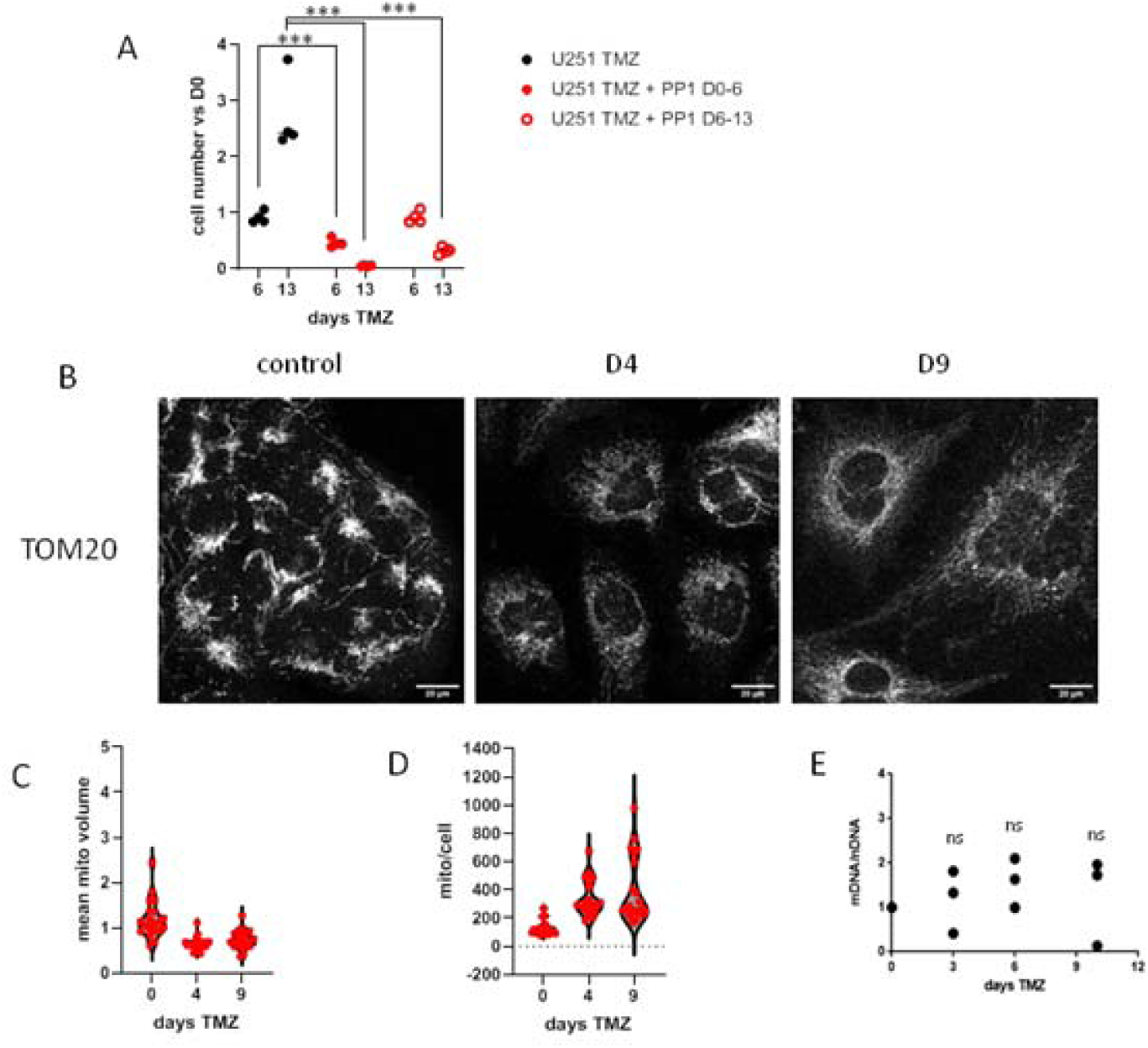
A. PP1 was added to U251 during TMZ treatment for the time indicated on the graph. Cells were counted by flow cytometry and normalized to the cell count at D0 (***: p<0.001). B. The morphology of mitochondria was observed in TMZ-treated U251 EGFR-cells by the staining of cells with a TOM20 antibody (scale bar: 20µm). C. The mean mitochondria volume (C) and number of mitochondria (D) was estimated by the Fiji software (Mitochondria Analyzer plugin) and divided by the number of cells in each field. The analysis was performed from at least six images per time point. E. Total DNA was extracted from TMZ-treated U251 EGFR-cells and the ratio mtDNA/nDNA was calculated as described. The graph illustrates three independent experiments.

### EGFR and TBK1 expressions are prognostic factors in Src-low GBM

Given the role of Src-activated EGFR and TBK1 axis in TMZ resistance unravelled above, we explored the prognostic value of their expression in available datasets from GBM patients in the TCGA_GBM (HG-UG133A) database (528 samples), through the GlioVis portal (https://gliovis.bioinfo.cnio.es) (21). The median mRNA expression was used as a cutoff value for the definition of low- and high-expressing patients. No obvious correlation between GBM patients survival and either EGFR (figure 6A), Src (figure 6B) or TBK1 (figure 6C) mRNA expression alone was established. No positive or negative correlation was observed between the three mRNAs expressions (suppl fig 6). Nevertheless, when considering only the Src-low GBM population, we observed a significant difference between the four subgroups identified according to EGFR and TBK1 stratification (figure 6D) (log rank p value = 0.0389), with a better prognosis for the “triple low” subgroup (EGFR-low/Src-low/TBK1-low). The median survival of this “triple low” subgroup (median=15.8 months, n=60) was significantly longer than the other Src-low subgroups (median=12.7 months, n=210) (figure 6E). These data advocate that enhanced expression of either EGFR and/or TBK1 worsens prognosis when Src itself is limiting, and is consistent with their pro-tumoral role downstream of Src.

**Figure 6:**
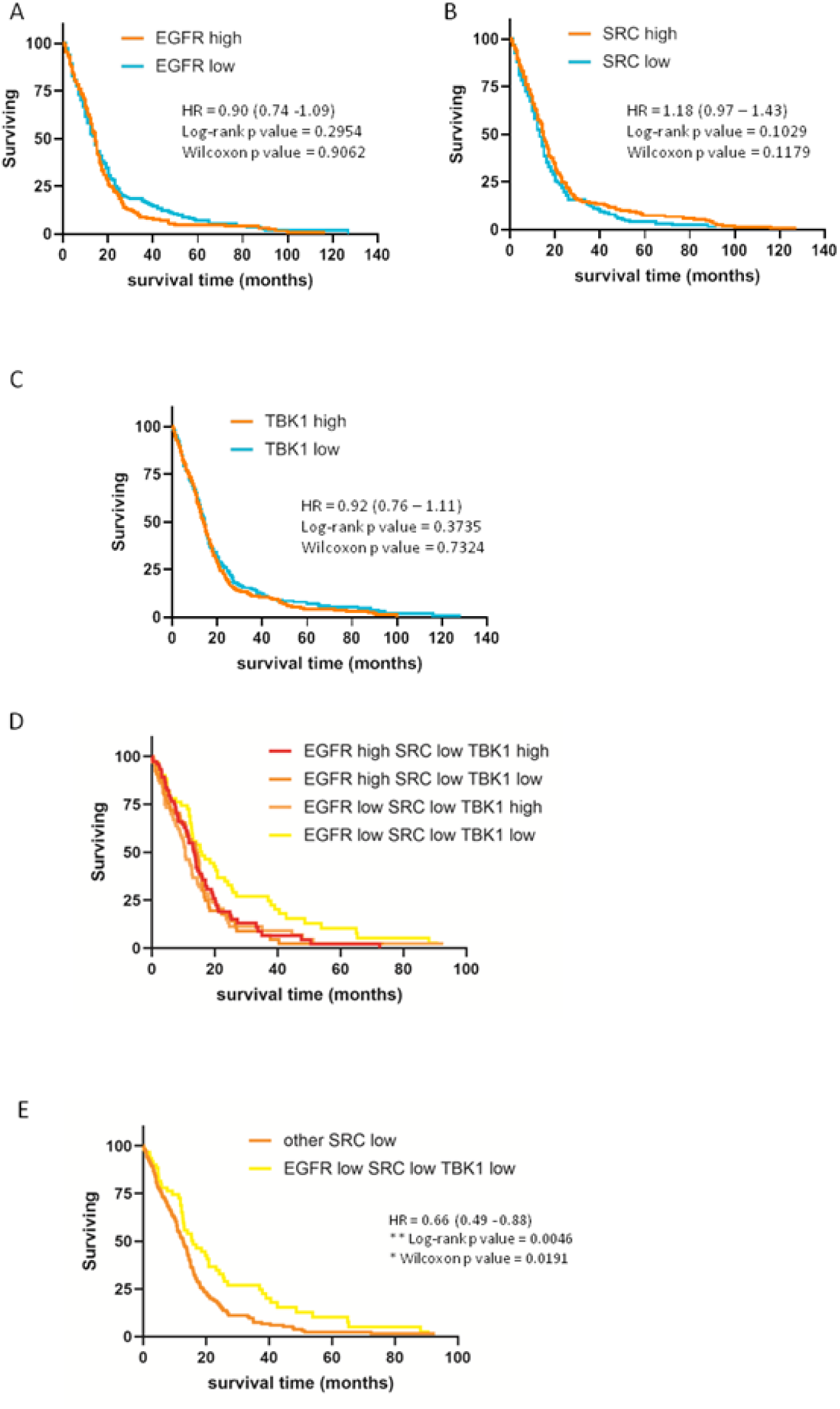
A. mRNA analyses from the TCGA_GBM (HG-UG133A) database to stratify patients according to EGFR expression (cutoff = median). The survival of both groups is shown by the Kaplan-Meier curve obtained from data extracted from GlioVis (orange: EGFR-high; cyan: EGFR-low). The same analysis was performed with Src mRNA expression (B) and TBK1 (C). D. GBM patients from the TCGA_GBM dataset were stratified according to their combined EGFR/Src/TBK1 expression. The Kaplan-Meier curve of the Src-low subgroups patients is shown. E. Data from D were analyzed by comparing the EGFR-low/Src-low/TBK1-low patients to other Src-low patients.

### Mitophagy is induced in the recovery phase in a TBK1/EGFR-dependent way

Autophagy is both a death and a survival program and is often implicated in the stress-recovery processes. During the acquisition of TMZ resistance, autophagy was found to be transiently induced in U251 cells as shown by the activation of LC3 measured by its lipidation (figure 7A) or the formation of LC3 aggregates visible by microscopic observation (suppl fig 7a). We questioned the implication of autophagy in the process of mitochondrial quality control and in the recovery process from the intermediate, adapted mitochondria to the mitochondrial phenotype of TMZ-resistant cells. We used a fluorescent protein addressed to the mitochondrial inner membrane (mEmerald-Mito7) as a reporter for mitochondrial turnover. U251 cells were transfected and mEmerald-expressing cells were sorted by flow cytometry to ensure optimal expression. Cells were treated by TMZ and mEmerald expression was measured by flow cytometry from naive to TMZ-resistant cells (D21). We observed a significant decrease in the fluorescence from D6 on with a maximum around D13, and then the expression of mEmerald raised again to reach a level similar to that of cells grown in the absence of TMZ for the same time (figure 7B). To further document the occurrence of an autophagy-driven mitochondrial recycling process, we investigated the proximity between mitochondria and autophagic vesicles. A proximity ligation assay revealed the presence of LC3 at the surface of the mitochondrial membrane (represented by endogenous TOM20-LC3 PLA, suppl fig 7b) around D10 (figure 7C). Among proteins implicated in the autophagic process, our proteomic analysis highlighted some actors upregulated during the intermediate time points (suppl fig 7c). The autophagy receptors SQSTM1 (or p62) and CALCOCO2 (or NDP52) were significantly overexpressed from D6 to D10 vs D0 and from D3 to D13 vs D0 respectively (p<0.01). GABARAP and GABARPL2, members of the LC3/GABARAP family proteins, were also upregulated during TMZ resistance acquisition (GABARAP is overexpressed at D6 vs D0 p<0.01 and at D10 vs D0 p<0.05 and GABARAPL2 was overexpressed at D10 vs D0 p<0.01). SNX30, implicated in the autophagosome assembly, was also overexpressed at D3, D13 (p<0.05), D6 and D10 (p<0.01) vs D0. TBK1 is an early player of mitophagy since it has the ability to initiate the phagophore formation (22,23). Accordingly, proximity ligation assay revealed the proximity between TBK1 and the mitochondrial outer membrane around D6, short before LC3 recruitment to the mitochondria (figure 7C). In contrast, at D6, we failed to detect an increase (and instead noted a slight decrease) of TFAM-LC3 proximity (figure 7C) indicating that this interaction recently demonstrated as an alternative initiator of the autophagic degradation of mtDNA-TFAM complexes (24), may not occur in our model. We added 3-methyladenine (3-MA), an early inhibitor of autophagy, to mEmerald-Mito7 expressing cells during TMZ treatment, either before (D6-10) or after (D10-D13) observation of LC3 proximity to the mitochondrial membrane. Strikingly, 3-MA abolished the decrease in mitochondrial fluorescence at D13 when added before LC3 recruitment to mitochondria (suppl fig 7d). This suggests that the increase in the mitochondrial turnover observed during TMZ treatment is linked to an autophagic recycling of mitochondria. At the functional level, the early inhibition of autophagosome formation by 3-MA drastically impacted the population viability when applied around D6 (either D3 to D6 or D6 to D10) (figure 7D). When applied later, from D10 to D13, the proliferation of the TMZ-resistant population was significantly delayed but cells started proliferating when 3-MA was removed. Note that the number of viable cells at D20 when TMZ was applied alone was slightly lower than in other experiments due to cells overconfluence. Selective degradation of TBK1 by a specific PROTAC molecule between D6 and D10 drastically reduced the amount of TOM20-LC3 PLA spots at D10 (suppl fig 7e). EGFR upon ligand-independent activation activates TBK1 by a transient interaction (19). We also investigated its role in the recruitment of LC3 in the vicinity of mitochondria. EGFR-deficient cells showed no significant change in TOM20-LC3 or TOM20-TBK1 proximity induced by TMZ (figure 7E). Altogether, this establishes the role of both EGFR and TBK1 in the mitochondrial turnover during the acquisition of resistance.

**Figure 7:**
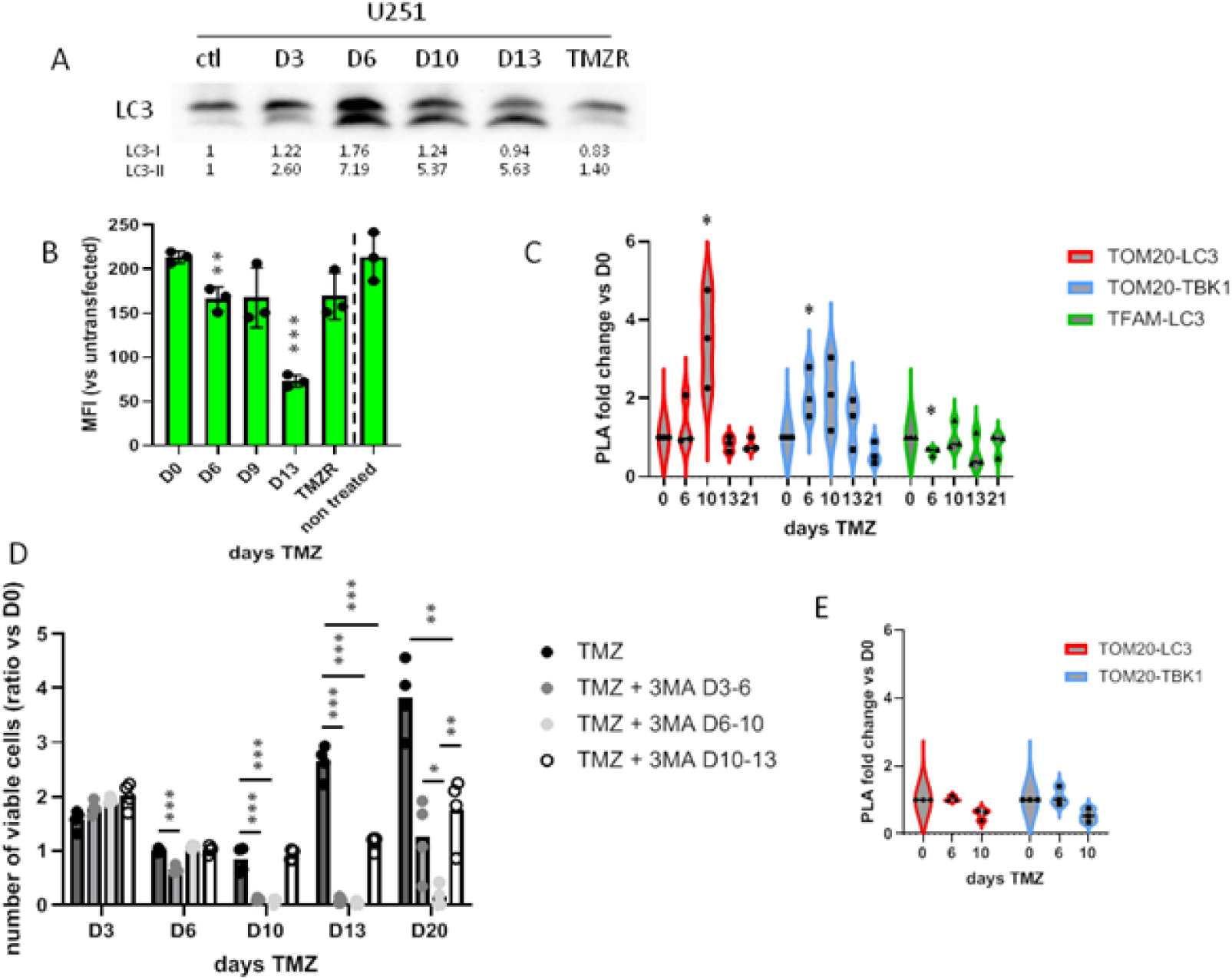
A. Protein lysates were obtained from TMZ-treated U251 and U251 EGFR-cells. LC3 was detected by Western Blot and signal was measured by ImageJ and normalized by the total protein staining intensity. The blot shown is representative from three independent experiments. B. U251 mEmerald-Mito7 were treated by TMZ and mEmerald fluorescence was measured at the indicated time point by flow cytometry. The graph shows the ratio of MFI of transfected cells to non-transfected cells at each timepoint. C. PLA was realized with the couple of antibodies indicated in TMZ-treated U251 cells. The number of spots was normalized to the number of spots at D0. (*: p<0.05 vs D0) D. 3MA was added to TMZ for the indicated time. Cells were counted by flow cytometry as described above and normalized to the cell count at D0. (**: p<0.01; ***: p<0.001). E. The same PLA experiment as in C was realized with TMZ-treated U251 EGFR-cells.

## Discussion

The acquisition of secondary chemoresistance is a complex and sequential phenomenon. In this study, we took advantage from a model of chemoresistance to TMZ in a GBM cell line described in Rabé *et al.* (8) and conducted a longitudinal, non-supervised global proteomic analysis. This longitudinal analysis strikingly highlights the prevalence of mitochondrial events during the resistance acquisition process (figure 1), which prompted us to focus on the mitochondrial adaptations required to the emergence of the drug-resistant population. GSEA analyses reveal the increase in the oxidative metabolism of cells during the intermediate states described, confirming the metabolic modifications observed by Rabé *et al*. (8). These modifications translate into a transient peak of mitochondrial ROS around D6 (figure 2). Of note, data obtained with respiration-incompetent cells (U251 Rho0) or NAC-treated U251 cells show that this ROS peak is hormetic in the acquisition of chemoresistance. Indeed, ROS both induce cell death and govern the emergence of the long-term drug resistant population, thereby highlighting their Janus-faced nature, as already described (5). Of note, the peak in the mitochondrial oxidative metabolism is not correlated with a massive increase in the mitochondrial mass or in the expression of proteins from the mitochondrial oxidative complexes (figure 2 and suppl fig 3). The metabolic adaptation may thus rather be due to a reorganization of the respiratory complexes into distinct supra-molecular complexes. The increase of mitochondrial Ca2+ linked to the raise in ER-mitochondria contacts may also participate in the increased activity of the respiratory oxidative complexes. The close apposition of neighboring membranes to mitochondria has been identified during early apoptotic stress, which may authorize the transfer of bioactive molecules between organelles, although membranes proximity is not sufficient by itself (15,25,26,27). An oscillation of mitochondrial Ca2+ was recently evidenced in a context of ER-mitochondria-plasma membrane platform, regulated by EGFR (27). In our model, we were directed to EGFR study by the fact that TBK1 is activated at the time of the ROS peak independent from the cGAS-STING pathway. The recruitment of TBK1, and subsequent phosphorylation, by the Src-activated EGFR has been previously demonstrated (10,19). The phosphorylation of EGFR at the Src target Y845 is the hallmark of mitochondrial EGFR translocation and the mitochondrial localization of EGFR induces metabolic modifications (29,30). EGFR Y845 is phosphorylated during TMZ treatment in a Src-dependent manner. Consistent with the role of EGFR demonstrated in other studies, in the absence of EGFR we observed an alteration in the mitochondrial modifications induced by TMZ in the early intermediate stages (mitochondrial fusion, mtDNA replication and transcription, ER-mitochondria contacts, TFAM distribution), suggesting that EGFR impacts on the mitochondrial adaptation. Of note, EGFR presence rather than its tyrosine kinase activity seems to be required. We observed that Src inhibition both impacts on the emergence of the TMZ-resistant population when applied early (D0-D6) or later (D6-D13). This suggests that mitochondrial EGFR may be required both to the stress induction program (sensitive to DTC transition) and during the stress resolution process (DTC to DRC transition). We measured the activation of Src independently from EGFR, yet we did not identify the activator of Src during TMZ treatment. Among the many potential proteins implicated in its activation (10,31,32,33), none was detected in our proteomic analysis or significantly deregulated by TMZ. We thus focused on the SH2- and SH3-binding proteins, and noticed that PTTG1 expression, a protein reported to prevent Src dephosphorylation induced by saracatinib (34), peaked at D6 in our proteomic analysis. Further studies will be required to establish if PTTG1 is actually implicated in Src activation during TMZ treatment.

During the resolution step, autophagy is induced and mitochondria are transiently degraded. We actually observed the recruitment of LC3 at the mitochondrial outer membrane, arguing for mitochondria-specific autophagy. The expression of a mitochondrial fluorescent protein, under the control of the strong CMV promoter (encoded by the mEmerald-Mito7 plasmid), unveiled an increase in the mitochondrial recycling, abolished by the early inhibition of autophagosome formation (3-MA). EGFR expression is required to the recruitment of LC3 to the mitochondrial outer membrane. This recruitment is preceded by the recruitment of TBK1 to the mitochondrial membrane and the targeted degradation of TBK1 (by PROTAC strategy) abolishes LC3 recruitment to the mitochondrial membrane. We thus propose that mitochondrial EGFR recruits TBK1 to the mitochondrial membrane, thereby inducing TBK1 activation (19), the subsequent recruitment of LC3 in the proximity of mitochondria. This might initiate autophagosome formation in itself, as it was shown that forced recruitment of Atg8-family proteins, including LC3 proteins, is sufficient to trigger mitophagy (35). TBK1 is a recognized regulator of mitophagy and contributes to the initiation of this mitochondrial quality control process by several means. Several studies report the interaction of TBK1 with the mitophagy receptors OPTN, NDP52 and/or p62 as a major regulatory point for the initiation of the autophagophore formation (36,37,38). Recent findings also reported the involvement of TBK1 in the autolysosome formation (39). We show here that mitochondrial, Src-activated EGFR might be responsible for TBK1 accumulation and auto-activation as initially described by Chakraborty *et al.* (19) for ligand-independent activated EGFR, and for the subsequent recycling of altered mitochondria by mitophagy. Notably, the apposition of ER to the mitochondrial outer membrane, as observed in our study, might provide the supply of lipids required for the autophagosome formation (40,41,42). We finally inhibited the early step of autophagy by the use of 3-MA and established the major role played by mitophagy initiation in the resolution of cell stress induced by TMZ (D6-10) and the emergence of the proliferation-active, TMZ-resistant cell population.

Although previous studies have described cellular modifications required for the appearance of the TMZ-resistant population in U251 glioblastoma cell line, our study clearly highlights the predominant role of mitochondrial adaptations in this process by a global, non-supervised proteomic approach, both during the early step of hormetic mitoROS peak induction and in the resolution of stress which is a prerequisite to the emergence of the proliferation active population. A recent large scale study combining genomics, transcriptomics and proteomics was recently conducted on a large cohort of primary and recurrent GBM (43). This study pointed out the interest of proteomics when combined with other techniques since information was only partially redundant between approaches. The authors identified several alterations shared by recurrent versus primary tumors. Interestingly, some of the alterations were recovered in our longitudinal proteomic analysis in the intermediate time points (D6-D10). This study also highlights a regulation of EGFR with an opposite correlation between transcriptomic and proteomic data (43), suggesting post-translational regulation. This finding is consistent with a specific degradation of mitochondria-associated EGFR by autophagy. Our proteomic data indicate a transient decrease in overall EGFR abundance during the transient stages (D6-D10), although the recruitment of LC3 to the mitochondria was not significantly detected until D10 in PLA experiments. Our proteomic analysis further focused on the role played by Src-activated EGFR and its partner TBK1 in the resolution of the hormetic ROS stress, enabling the emergence of the resistant and proliferative U251 population. Proximity ligation assays, genetic modifications and PROTAC experiments reveal the pivotal role of the Src-activated EGFR / TBK1 couple in the mitochondrial recycling autophagic process. The regulation of this pathway and its activation in other models of tumors expressing high levels of EGFR will be the subject of further investigations.

## Acknowledgements

We are grateful to Nicolas Bidère for the generous gift of PROTAC TBK1 and negative control.

We acknowledge the IBISA MicroPICell facility (Biogenouest), member of the national infrastructure France-Bioimaging supported by the French national research agency (ANR-10-INBS-04).

We also thank the Cytocell - Flow Cytometry and FACS core facilty (SFR Bonamy, BioCore, Inserm UMS 016, CNRS UAR 3556, Nantes, France) for its technical expertise and help, member of the Scientific Interest Group (GIS) Biogenouest and the Labex IGO program supported by the French National Research Agency (ANR-11-LABX-0016-01).

This project was supported by the Ligue Contre le Cancer Comités du Finistère et de Loire-Atlantique.

