## Supplementary data for "EGFR/TBK1-dependent mitochondrial quality control contributes to acquired resistance to temozolomide"

**Supplementary figure 1**: a) U251 cells were treated by TMZ for the indicated time and adherent and floating cells were collected. Cells were pelleted by centrifugation and stained by ToPro3 for 10mn at room temperature. The percentage of ToPro3 positive cells was counted by flow cytometry. The graph represents the mean +/- sd of three independent experiments. b) PCA was performed from the proteomic data from 5 independent experiments as described in the methods section (Spectronaut® statistical tools). c) The list of proteins significantly upregulated at D6 vs D0 was submitted to EnrichR for GSEA analysis. The 10 top pathways identified by Gene Ontology Biological Process (up) and Gene Ontology Cell Component (bottom) are represented on the bubble graphs as indicated in the material and methods section. d) The list of proteins significantly upregulated (up) or downregulated (bottom) at D21 vs D0 was submitted to EnrichR for GSEA analysis. The 10 top pathways identified by Gene Ontology Biological Process are shown. e) The list of proteins significantly upregulated at D6 vs D0 and downregulated at D21 vs D0 (up) or the inverse (bottom) was submitted to EnrichR for GSEA analysis. The 10 top pathways identified by Gene Ontology Biological Process are shown.

a

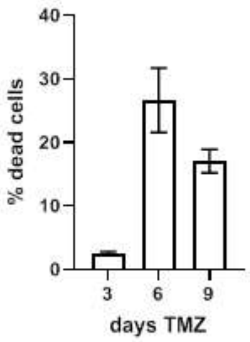


b

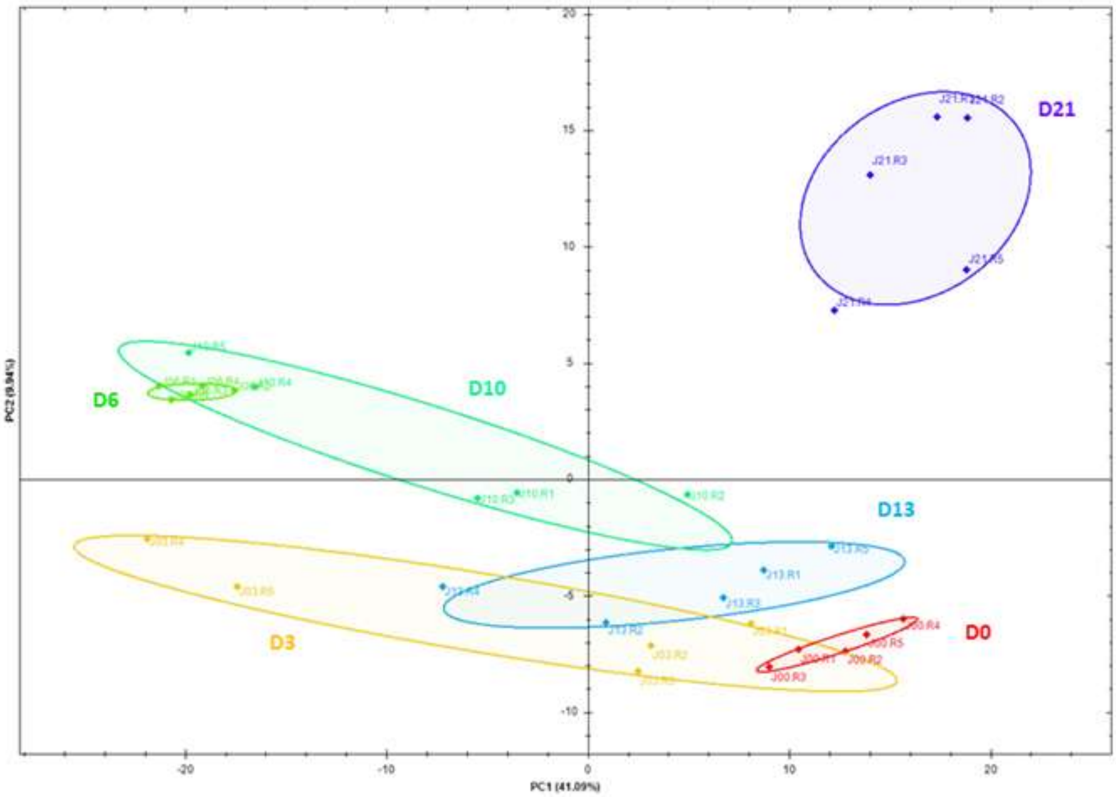

c Up D6 vs D0

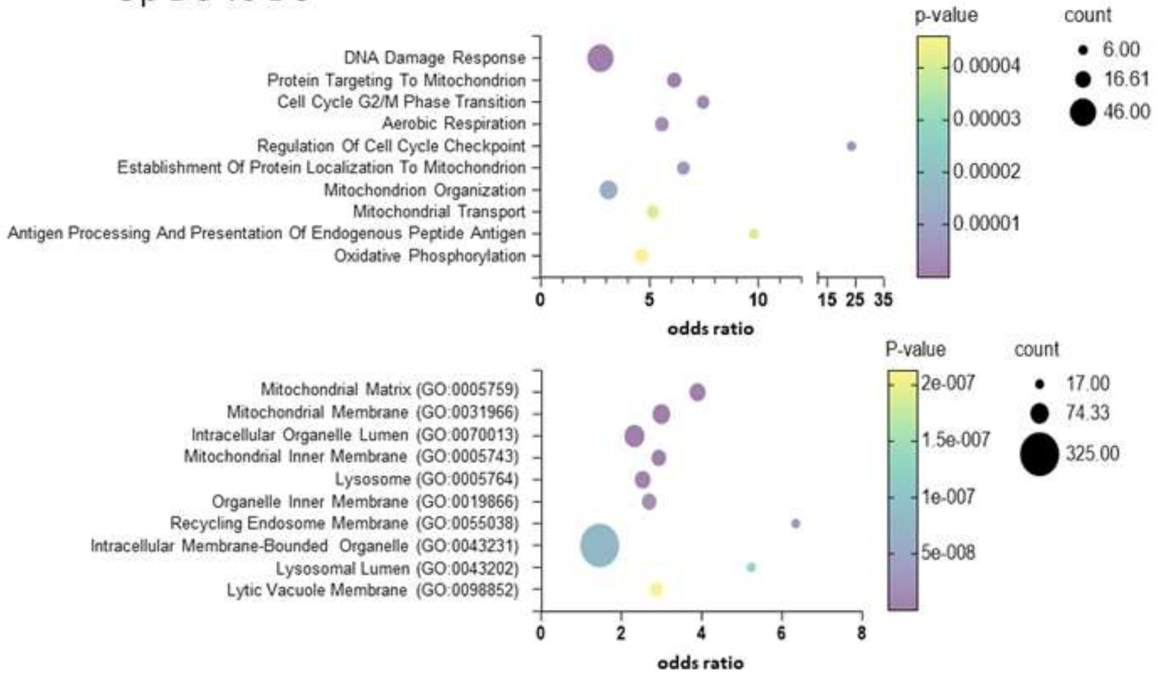


d GO biological process

Up D21 vs D0

Down D21 vs D0

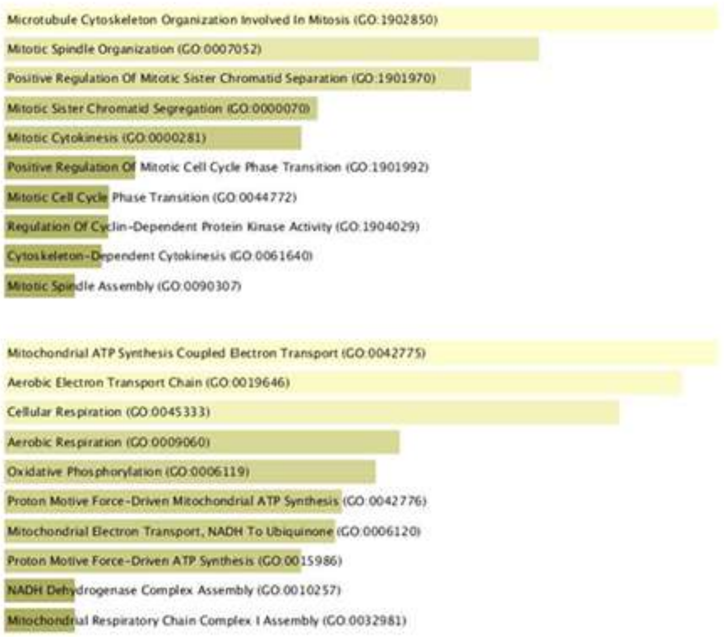

e

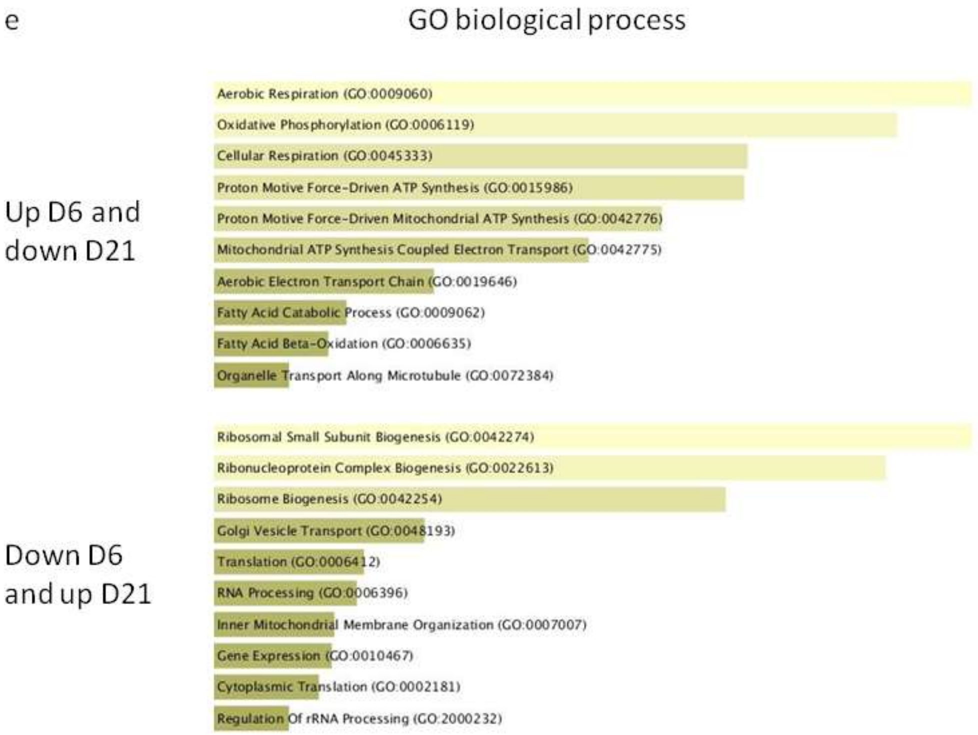

GO biological process
Up D6 and down D21
Aerobic Respiration (GO:0009060)
Oxidative Phosphorylation (GO:0006119)
Cellular Respiration (GO:0045333)
Proton Motive Force-Driven ATP Synthesis (GO:0015986)
Proton Motive Force-Driven Mitochondrial ATP Synthesis (GO:0042776)
Mitochondrial ATP Synthesis Coupled Electron Transport (GO:0042775)
Aerobic Electron Transport Chain (GO:0019646)
Fatty Acid Catabolic Process (GO:0009062)
Fatty Acid Beta-Oxidation (GO:0006635)
Organelle Transport Along Microtubule (GO:0072384)
Down D6 and up D21
Ribosomal Small Subunit Biogenesis (GO:0042274)
Ribonucleoprotein Complex Biogenesis (GO:0022613)
Ribosome Biogenesis (GO:0042254)
Golgi Vesicle Transport (GO:0048193)
Translation (GO:0006412)
RNA Processing (GO:0006396)
Inner Mitochondrial Membrane Organization (GO:0007007)
Gene Expression (GO:0010467)
Cytoplasmic Translation (GO:0002181)
Regulation Of rRNA Processing (GO:2000232)

**Supplementary figure 2**: a) Mitochondrial morphology was assessed as indicated in Figure 2A at Day 12 of TMZ treatment. The image shown is representative of three independent experiments (scale bar: 20µm). b) The log2FC values of candidates differentially expressed and involved in mitochondrial fission (left) and fission (right) according to MitoCarta3.0 were calculated for each of the five replicates and the mean of the five values is represented on the graph. The results of the paired Student t-tests are described in the main text. c) and d) RT-QPCR was performed on total mRNA extracted from TMZ-treated U251 cells. RPLPO was used as housekeeping gene and non-treated U251 cells were used as reference. The graph illustrates data from three independent experiments (*: p<0.05, **: p<0.01). e) The log2FC values of candidates differentially expressed and involved in mitochondrial DNA replication (left) and transcription (right) according to MitoCarta3.0 were calculated for each of the five replicates and the mean of the five values is represented on the graph. The results of the paired Student t-tests are described in the main text.

a

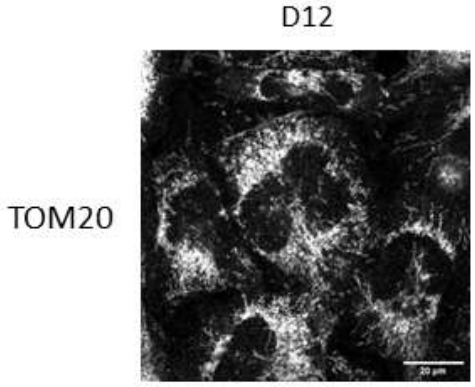


b

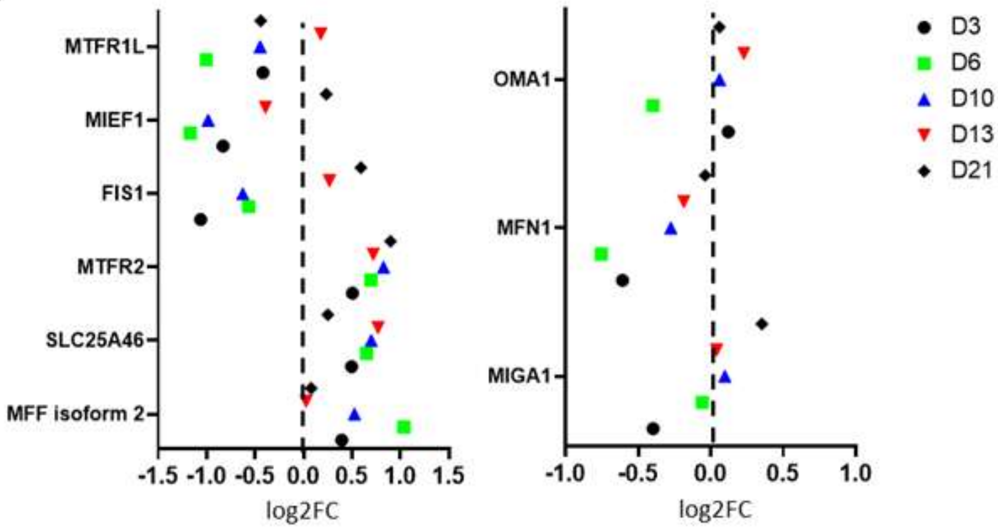

c

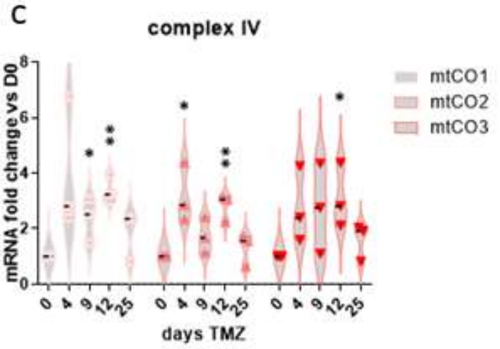
complex IV
mRNA fold change vs D0
0
2
4
6
8
mtCO1
mtCO2
mtCO3
0 4 9 12 25 0 4 9 12 25 0 4 9 12 25
days TMZ

d

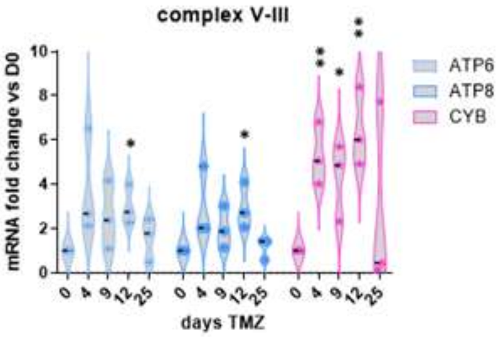
complex V-III
mRNA fold change vs D0
0
2
4
6
8
10
ATP6
ATP8
CYB
0 4 9 12 25 0 4 9 12 25 0 4 9 12 25
days TMZ

e

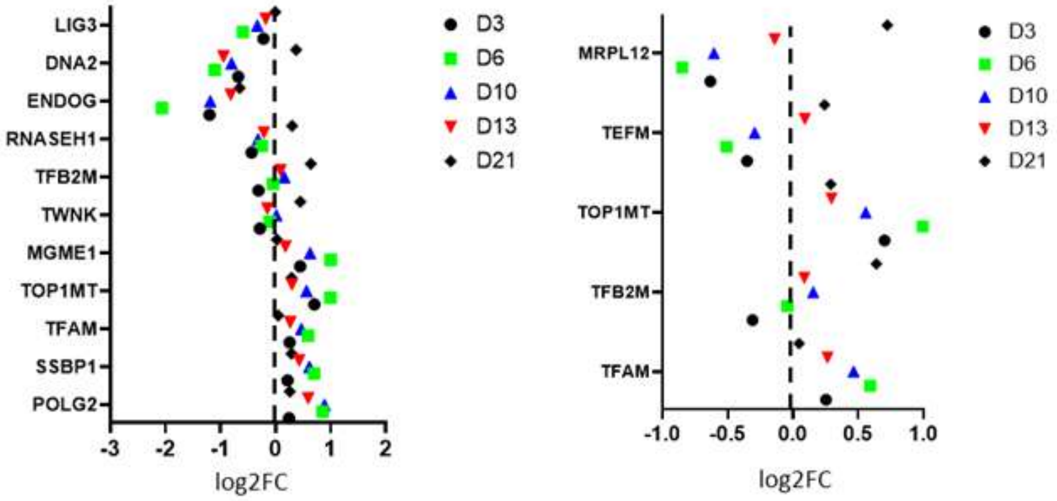
LIG3
DNA2
ENDOG
RNASEH1
TFB2M
TWNK
MGME1
TOP1MT
TFAM
SSBP1
POLG2
-3 -2 -1 0 1 2
log2FC
D3
D6
D10
D13
D21
MRPL12
TEFM
TOP1MT
TFB2M
TFAM
-1.0 -0.5 0.0 0.5 1.0
log2FC
D3
D6
D10
D13
D21

**Supplementary figure 3**: a) The log2FC of the proteins from the mitochondrial respiratory complexes was calculated for each replicate. Heat maps were drawn by the Prism software for each complex (complex I: CI to complex V: CV). b) U251 mEmMito7 cells were grown on glass coverslips and treated by TMZ for 6 days. Cells were stained by Rhod2AM as described in material and methods, then fixed and mounted with Prolog Gold with DAPI before confocal microscopic observation. Rhod2AM was essentially colocalized with mitochondria at D0 and D6 as shown on the images (left) and on the magnified inserts (right). The magnified area is shown by the white frame on the left images. Scale bar: 20µm. c) U251 cells were grown on glass coverslips in 24-well plates and treated by TMZ for the indicated time. Cells were fixed and a proximity ligation assay (PLA) was performed with the VDAC-IP3R couple of antibodies to visualize mitochondria-ER contacts according to the manufacturer instructions. Primary antibodies were omitted for negative control. Prolong Gold with DAPI was used for coverslips mounting (scale bar: 20µm). d) U251 and U251 Rho0 cells were treated by TMZ for 72h. TMZ-treated and untreated cells were stained by MitoSOX Red (1µM, 20mn). Cells were then collected and analyzed by flow cytometry on a BD Accuri C6 instrument. The graph represents the mean value of FL2med from three independent experiments. e) VDAC-IP3R PLA was performed on TMZ-treated U251 cells as described in Figure 3H but the number of spots per cell was normalized to the number of spots per cell in the cells without NAC at the same time point. (**: $p<0.01$; ***: $p<0.001$ Student t-test vs no NAC).

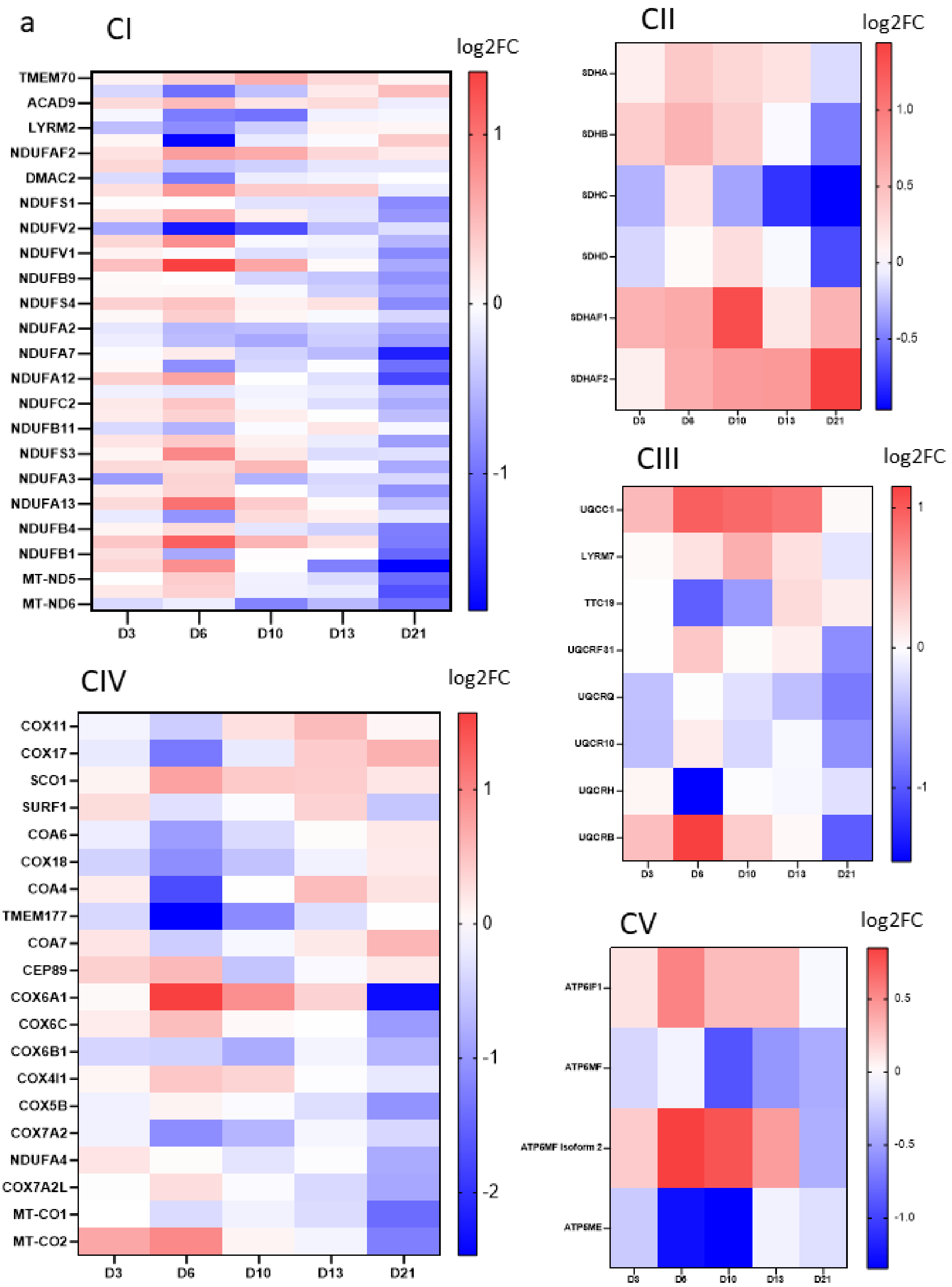

a
CI
log2FC
TMEM70
ACAD9
LYRM2
NDUFAF2
DMAC2
NDUFS1
NDUFV2
NDUFV1
NDUFB9
NDUFS4
NDUFA2
NDUFA7
NDUFA12
NDUFC2
NDUFB11
NDUFS3
NDUFA3
NDUFA13
NDUFB4
NDUFB1
MT-ND5
MT-ND6
D3
D6
D10
D13
D21
1
0
-1
CII
log2FC
SDHA
SDHB
SDHC
SDHD
SDHAF1
SDHAF2
D3
D6
D10
D13
D21
1.0
0.5
0
-0.5
CIII
log2FC
UQCC1
LYRM7
TTC19
UQCRFS1
UQCRQ
UQCR10
UQCRH
UQCRB
D3
D6
D10
D13
D21
1
0
-1
CIV
log2FC
COX11
COX17
SCO1
SURF1
COA6
COX18
COA4
TMEM177
COA7
CEP89
COX6A1
COX6C
COX6B1
COX4I1
COX5B
COX7A2
NDUFA4
COX7A2L
MT-CO1
MT-CO2
D3
D6
D10
D13
D21
1
0
-1
-2
CV
log2FC
ATP5IF1
ATP5MF
ATP5MF Isoform 2
ATP5ME
D3
D6
D10
D13
D21
0.5
0
-0.5
-1.0

b

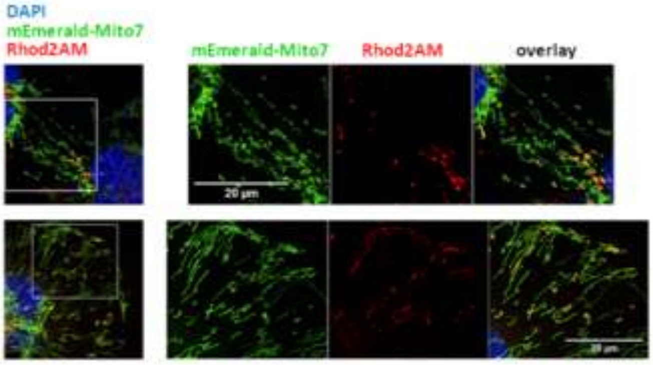
DAPI
mEmerald-Mito7
Rhod2AM
mEmerald-Mito7
Rhod2AM
overlay
D0
D6
20 µm

c

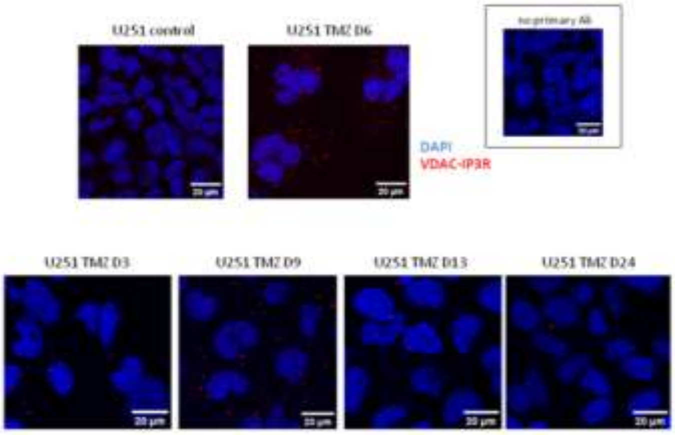
U251 control
U251 TMZ D6
no primary Ab
DAPI
VDAC-IP3R
U251 TMZ D3
U251 TMZ D9
U251 TMZ D13
U251 TMZ D24
20 µm

d

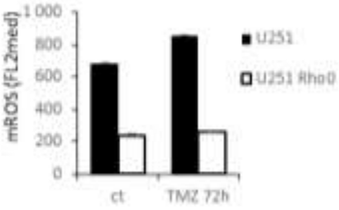
mROS (FL2med)
1 000
800
600
400
200
0
ct
TMZ 72h
U251
U251 Rho0

e

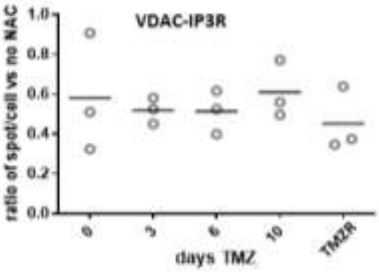
VDAC-IP3R
ratio of spot/cell vs no NAC
1.0
0.8
0.6
0.4
0.2
0.0
0
3
6
10
TMZR
days TMZ

**Supplementary figure 4**: a) CRISPR Cas9-induced knock-down of cGAS or STING was performed on U251 cells by the use of the lentiviral vectors described in Lohard *et al.* (Nat. Commun., 2020). Control cells consist of cells treated by a non-targeting gRNA. Cells were seeded in 24-well plates and treated by TMZ from the next day. At the indicated time point, cells were collected and counted and the number of cells was normalized by the cell number at D0 for each replicate. b) The cells described in a) were treated by TMZ for the indicated time points and cells were fixed, permeabilized and stained with a pTBK1 antibody coupled to a fluorescent secondary antibody. Cell fluorescence was analyzed by flow cytometry and the median fluorescence was normalized by the mean fluorescence of control cells at each time point. c) U251 were treated for 6 days by TMZ 50µM in the absence or presence or PP1 10µM, a selective inhibitor of Src, or saracatinib 0.2µM, a dual inhibitor of Bcr-Abl and Src. Cells were collected, fixed, permeabilized and stained with an anti pSrc antibody then coupled to a fluorescent secondary antibody as described in the material and methods section. Cell fluorescence was analyzed by flow cytometry. Fluorescence intensity is expressed as the fold change vs the mean fluorescence measured in non treated (D0) cells (*: $p<0.05$). d) U251 cells were treated by TMZ or TMZ + PP1 (10µM) for 6 days. Cells were collected, fixed, permeabilized and stained with an antibody raised against total EGFR then coupled to a fluorescent secondary antibody. Cell fluorescence was analyzed by flow cytometry.

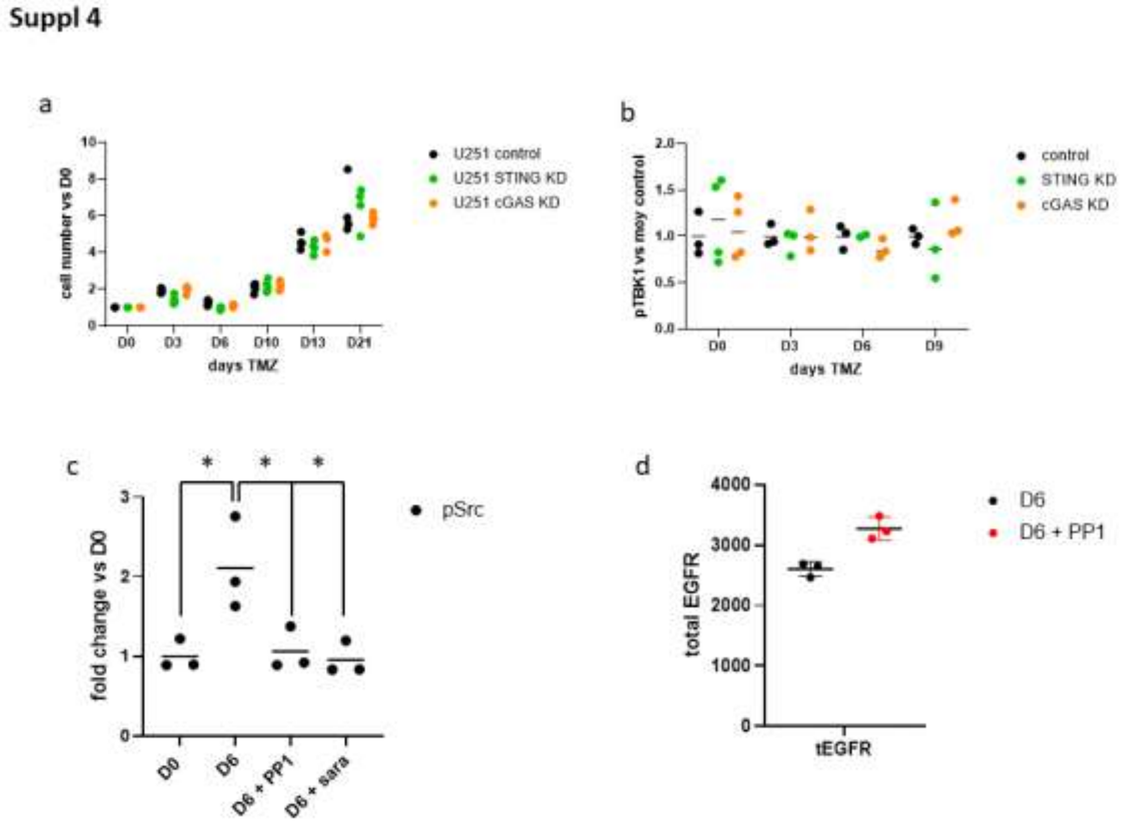

**Supplementary figure 5**: a) U251 cells were treated by TMZ 50µM with saracatinib 0.2µM added either between D0 and D6 or between D6 and D13. Cells were counted by flow cytometry and normalized to the cell count at D0. (***: $p<0.001$, *: $p<0.05$). b) U251 EGFR- cells were grown on glass coverslips and treated by TMZ for the indicated time. Cells were fixed and VDAC-IP3R PLA analysis was performed as described in the material and methods section. The number of spots per cell was normalized to the number of spots at D0 in each experiment. The graph is representative of three independent experiments. c) RT-QPCR was performed on mRNA extracted from U251 EGFR- cells as described in the main text. RPLPO was used as housekeeping gene and non-treated U251 cells were used as reference. The graph illustrates data from three independent experiments. d) U251 and U251 EGFR- cells were grown on glass coverslips, treated by TMZ and stained for TFAM at the indicated time points. Coverslips were mounted with Prolong Gold with DAPI and observed by confocal microscopy.

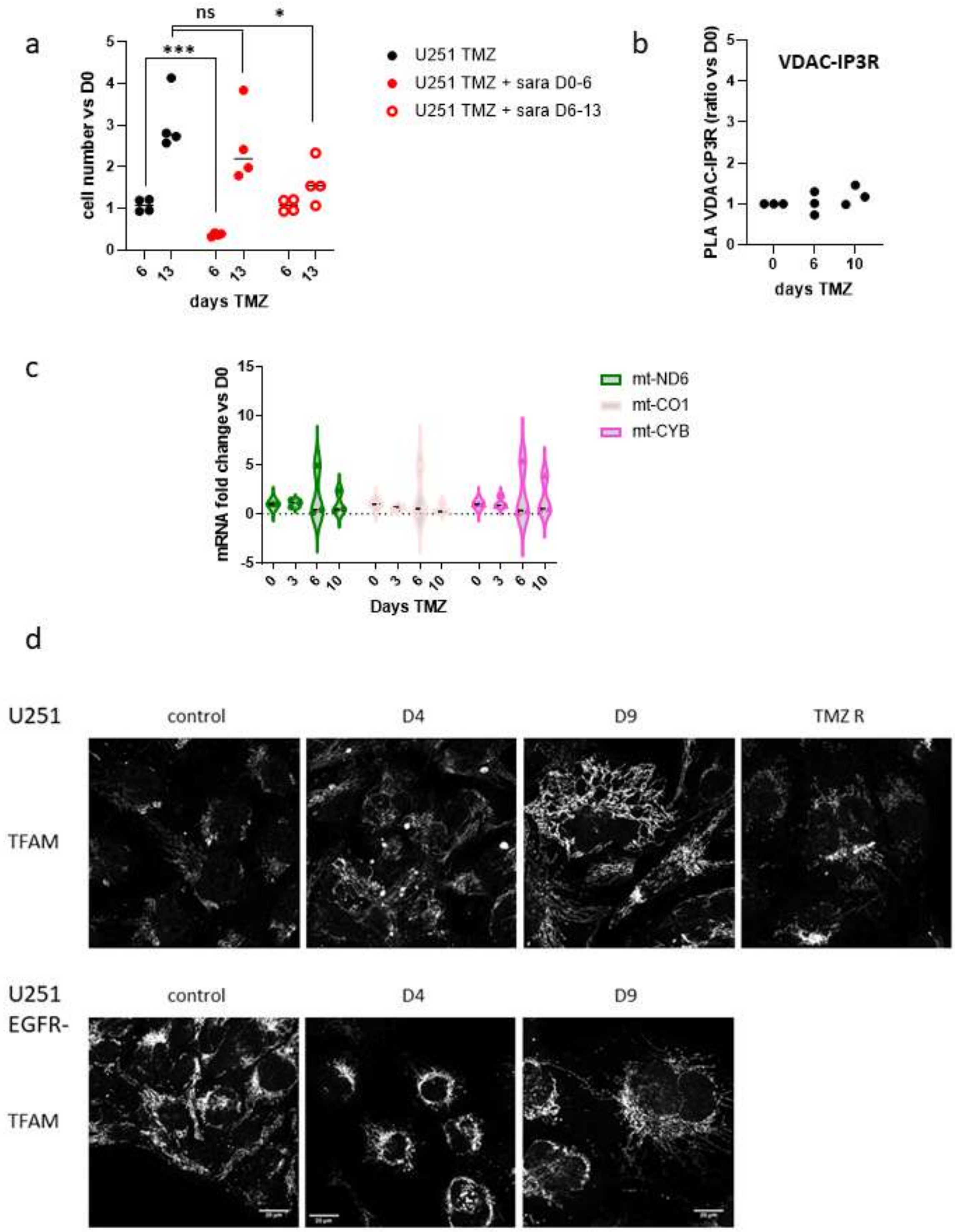
a
ns
*
***
cell number vs D0
0
1
2
3
4
5
6
13
6
13
6
13
days TMZ
U251 TMZ
U251 TMZ + sara D0-6
U251 TMZ + sara D6-13
b
VDAC-IP3R
PLA VDAC-IP3R (ratio vs D0)
0
1
2
3
4
5
0
6
10
days TMZ
c
mRNA fold change vs D0
-5
0
5
10
15
0
3
6
10
0
3
6
10
0
3
6
10
Days TMZ
mt-ND6
mt-CO1
mt-CYB
d
U251
control
D4
D9
TMZ R
TFAM
U251
EGFR-
control
D4
D9
TFAM

**Supplementary figure 6**: mRNA expression of EGFR, Src and TBK1 were obtained from the TCGA_GBM (HG-UG133A) dataset (528 samples). The correlation between EGFR and Src (a), EGFR and TBK1 (b) and Src and TBK1 (c) is shown on the graphs.

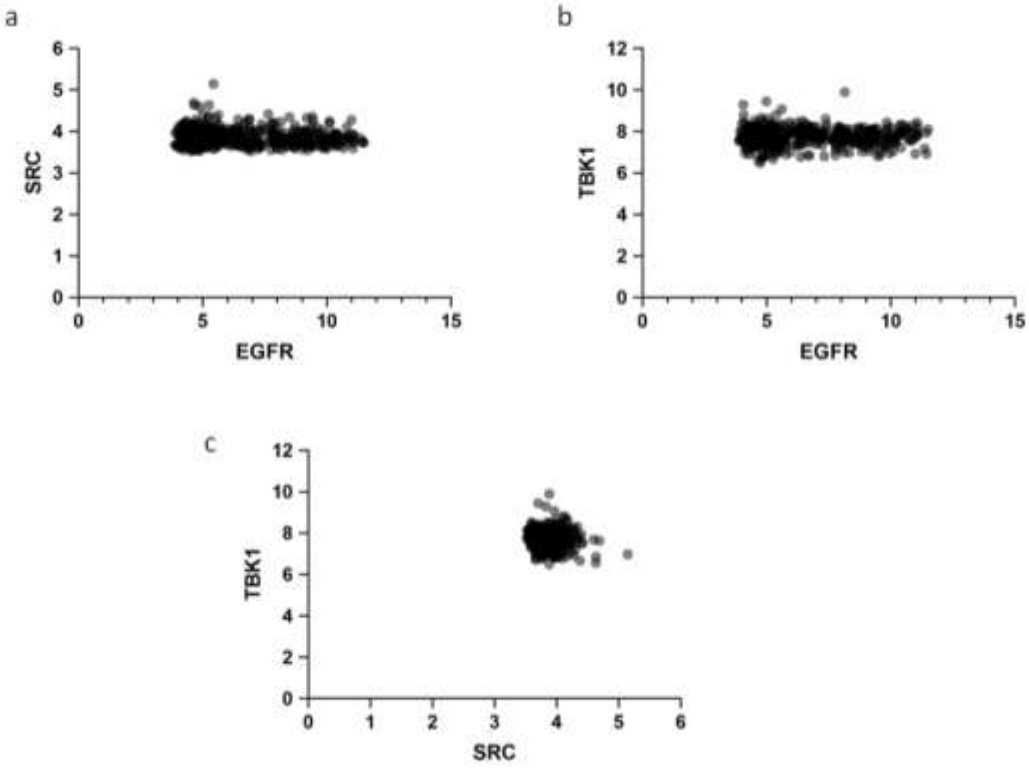

**Supplementary figure 7**: a) U251 cells grown on glass coverslips were treated by TMZ for the indicated time. Cells were fixed, permeabilized and stained by an antibody raised against LC3 (Cell Signaling Technology, #83506) followed by fluorescent secondary antibody. Cells were observed by confocal microscopy. The experiment was repeated 3 times. b) Representative images of the PLA performed for TOM20-LC3 proximity are shown. The insert shows an image of the PLA negative control. c) The log2FC values of candidates differentially expressed and involved in the autophagic process were calculated for each of the five replicates and the mean of the five values is represented on the graph. The results of the paired Student t-tests are described in the main text. d) U251-mEmerald-Mito7 cells were treated by TMZ for 13 days and 3-MA was added either before (D6-10) or after D10 (D10-D13). mEmerald fluorescence was measured by flow cytometry and MFI was normalized as described in the material and methods section. e) TOM20-LC3 PLA analysis was performed on U251 cells treated by TMZ for 10 days, treated by a TBK1 PROTAC or a negative control (100nM) added every day from D6 to D10. TBK1 proteolysis induced by PROTAC was assessed by western blot and one representative blot is shown above the graph (* indicates a non specific band). Three independent experiments were realized and the number of spots per cell was normalized to the number of spots in the control cells for each experiment. The insert on the right shows a representative image of the PLA TOM20-LC3 in U251 cells treated by the PROTAC TBK1. The scale bar for all microscopic images is 20µm.

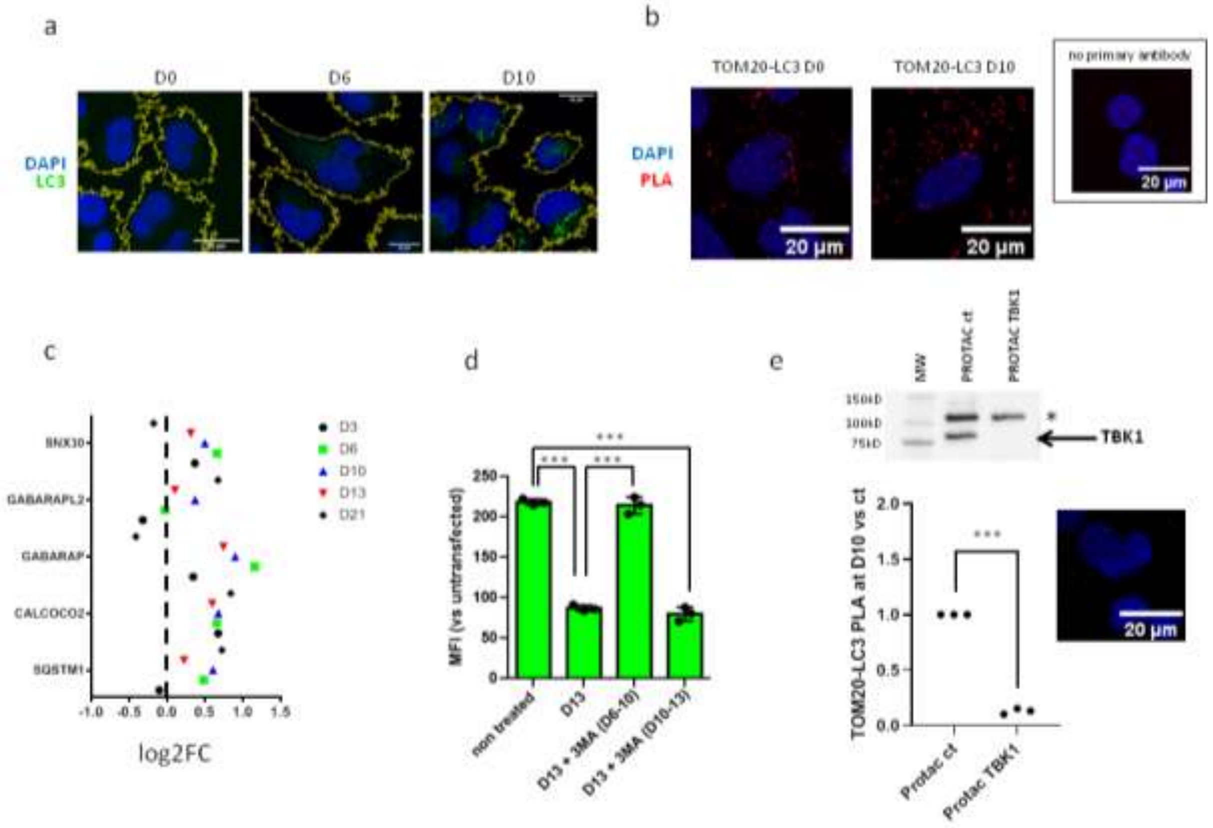
